# Spatial Transcriptomics Reveals Age-Specific Cancer Hallmark Architecture in Pediatric and Adult Brain Tumors

**DOI:** 10.64898/2026.11.13.766471

**Authors:** Noah T. Moffat, Clark T. Tolbert, Carter W. Middleton, Mustafa Sibai, Sergi Cervilla, Eduard Porta-Pardo, Matthew H. Bailey

## Abstract

Pediatric cancers are fundamentally different from their adult counterparts. Specifically, pediatric cancers carry fewer mutations, interact with developing immune systems, and have active developmental microenvironments. The tumor microenvironment (TME), which includes the surrounding stromal cells, immune cells, blood vessels, and extracellular matrix, plays a crucial role in cancer progression, metastasis, behavior, and treatment response. To better understand the effects of the pediatric tumor microenvironment on tumor gene expression, we acquired spatial transcriptomic data from 17 pediatric brain tumor samples and analyzed it through a Cancer Hallmarks framework. Cancer Hallmarks, as outlined by Hanahan and Weinberg, provide a comprehensive set of 14 biological capabilities that cancers acquire to enable tumor growth and metastasis. We compared the spatial gene expression patterns of the Cancer Hallmarks gene sets between the TME and neoplastic compartments in pediatric brain tumors. Compared with a similar study of adult brain tumors, in which Cancer Hallmark expression patterns clearly separated the TME and neoplastic compartments, we identified three hallmarks (“Deregulating Cellular Energetics,” “Senescent Cells,” and “Evading Growth Suppressors”) whose predominant cellular compartments differed between pediatric and adult tumors. In pediatric samples, these hallmarks were associated primarily with the TME rather than the neoplastic compartment. This shift implicates an age-specific rewiring of the tumor ecosystem. We also provide clear evidence of strong transcriptomic regulation in genes shared by pediatric and adult neoplastic compartments. Finally, we show how measures of spatial autocorrelation may be used to distinguish tumor grade.

**Significance:** Distinguishing between pediatric and adult tumors is essential for tailoring treatments to specific age groups and discovering new interactions between the tumor microenvironment (TME) and neoplasms. Pediatric brain tumors exhibit distinct biological features compared to their adult counterparts, including lower mutational burden and unique immune responses. However, whether these differences manifest primarily in tumors, the TME, or both has been unclear. In this study, we systematically compare the Cancer Hallmarks architecture using existing spatial transcriptomic gene sets and ask whether each set is more frequently observed in the neoplastic compartment versus the TME of pediatric tumors. By preserving the spatial resolution of gene expression, our findings elucidate pediatric tumor pathways, genetic factors, and how these interact with surrounding tissue. This framework may clarify how age-associated developmental programs shape growth-promoting states in pediatric tumors and their TME.

## Introduction

Brain tumors are the leading cause of cancer-related deaths in children (Patel et al. 2021). While treatments are often lifesaving, survivors of these cancers are highly likely to develop chronic health issues later in life (Patel et al. 2021). While our molecular understanding of pediatric brain tumors continues to improve, low mutation rates and developmentally primed microenvironments present research challenges where the majority of tumor signal is derived from very few molecular changes, unlike their adult counterparts. With new technologies, such as spatial transcriptomics, we can begin to assess interactions between tumors and their microenvironments while preserving spatial relationships. Characterizing the pediatric brain TME and neoplastic interactions is essential to understanding their roles in tumor progression and treatment response.

The Cancer Hallmarks framework provides a high-level approach to characterizing cancer by defining the essential traits that tumors acquire during tumorigenesis and progression (D. Hanahan and Weinberg 2000; Douglas Hanahan and Weinberg 2011; Douglas Hanahan 2022). These 14 Hallmarks represent fundamental biological capabilities that enable cancers to proliferate uncontrollably, evade growth suppressors, resist cell death, and metastasize, among other traits. A comprehensive evaluation of the spatial distribution of pediatric Cancer Hallmarks is of great importance, as their location and presentation may influence disease progression and therapeutic response. Moreover, many cancer treatments relate to the Cancer Hallmarks framework, e.g., antiangiogenics, immunotherapies, chemotherapy that induces genomic stability, and targeted therapies designed to disrupt proliferative cell signaling.

Recently, Sibai et al. 2025 leveraged the Cancer Hallmark structure to spatially resolve neoplastic and TME compartments in adult tumors (Sibai et al. 2025). Briefly, this study utilized spatial gene expression data to produce “hallmark scores” per spatial region through differential expression analysis of hallmark-associated genes. Importantly, Sibai et al. demonstrated that seven Cancer Hallmarks reach higher expression scores in adult neoplastic compartments, including “Resisting Cell Death,” “Senescent Cells,” “Evading Growth Suppressors,” “Deregulating Cellular Energetics,” “Nonmutational Epigenetic Reprogramming,” “Enabling Replicative Immortality,” and “Genome Instability and Mutation” (**Fig. 1A**, adapted from Sibai et al. 2025). The remaining six Cancer Hallmarks were found to have higher scores in the TME. Thus, the tumor and non-tumor compartments contributed evenly to the ecological dynamics of a pro-tumor environment. Of note, the Hallmark “Polymorphic Microbiomes” was excluded from their analysis because the assay was designed to capture human genetic material. Consequently, the analysis encompassed 13 of the 14 Cancer Hallmarks. Additionally, using clinical trial data from the DUTRENEO study, Sibai et al. found that spatial relationships of Cancer Hallmarks predicted patient response to immunotherapy. Given the predictive value of spatial Hallmark activity in adult tumors, applying a similar framework to pediatric brain tumors may uncover parallel mechanisms or reveal unique therapeutic opportunities.

**Figure 1:**
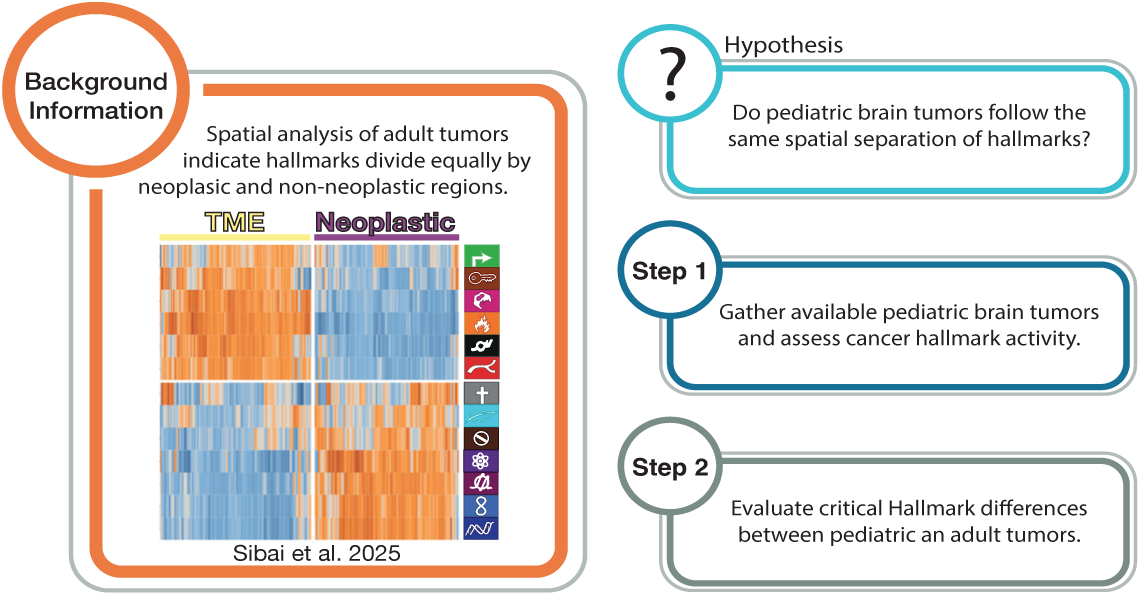
Background, questions, and overview of manuscript. Four boxes highlight background, question, and basic outline of the manuscript.

As such, we applied the spatial Cancer Hallmarks framework (Sibai et al. 2025) to study cancers in pediatric patients. Because the most common forms of pediatric cancer affect the central nervous system, we decided to study pediatric brain cancers. Here, we address the question: Do pediatric brain tumors follow the same spatial separation of Cancer Hallmarks as adult tumors? If not, which Hallmarks are more prominent in the TME or neoplastic compartments, and can we provide molecular insights into the structure and function of pediatric brain tumors (**Fig. 1B**)?

## Results

### ACTIVITY OF CANCER HALLMARKS IN PEDIATRIC BRAIN CANCERS FOLLOWS A SPATIALLY ORGANIZED ARCHITECTURE

We acquired spatial transcriptomic data from 17 untreated human pediatric brain tumor samples from two cancer types: pediatric ependymoma (n = 13, Fu et al., 2023) (Fu et al. 2023) and diffuse midline glioma (n = 4, Ren et al., 2023) (Ren et al. 2023). After inspecting each sample’s 10X Visium results (a spatial transcriptomics assay capable of capturing 5000 unique regions, i.e., spots, on a 6.5mm x 6.5mm tissue section), we determined that all of our data were of sufficient quality (**Table S1**). Overall, we observed 39,766 unique spots under tissue with an average of 23,378 genes detected per sample. Across all samples, the median number of genes detected per spot ranged from 2,751 to 7,221, with a median of approximately 3,742 genes per spot.

Spatial transcriptomic data were generated using the 10X Genomics Visium platform. Because each ∼55µm spot generated by Visium can contain several cells, we approximated single-cell resolution by subdividing each spot into six “subspots” of ∼20µm using BayesSpace (Zhao et al. 2021). We annotated each subspot implementing ESTIMATE tumor purity testing, clustering the results into 5 classifications ranging from neoplastic (1) to TME (5) (Yoshihara et al. 2013) (**Samples and Methods**). For downstream analysis, we assigned clusters 1 and 2 as the neoplastic compartment, 4 and 5 as the TME compartment, and cluster 3 as an intermediate buffer between neoplastic and TME compartments. Importantly, a sample’s neoplastic and TME compartments include all subspots annotated as “neoplastic” or “TME,” respectively, regardless of whether they are spatially contiguous.

Next, we measured the transcriptional activity of each of the 13 Cancer Hallmarks by generating “hallmark scores” for each subspot using a methodology nearly identical to Sibai et al. (**Samples and Methods**). Using unsupervised clustering methods, we clustered average hallmark scores for each sample’s neoplastic and TME compartments. We observed substantial differences in average hallmark scores between the two compartments (Neoplastic or TME, p-value < 2.2×10^−16^, Fisher’s exact test) and this clustering was not likely attributable to cancer type (p-value = 0.8, Mann-Whitney *U* test). Thus, like their adult counterparts, the shared attributes of the Neoplastic and TME compartments play a strong role in influencing spatial Hallmark patterns. Notably, our clustering of pediatric ependymoma and midline glioma samples identified three Cancer Hallmarks that differed from the adult clustering of spatial Cancer Hallmarks – “Deregulating Cellular Energetics,” “Senescent Cells,” and “Evading Growth Suppressors”. These three Hallmark scores, strongly expressed in the adult neoplastic compartment, were now strongly expressed in the pediatric TME compartment (**Fig. 2A**). The other 10 pediatric Hallmarks remained identical to the adult classification. Furthermore, four Cancer Hallmarks displayed higher sample-level heterogeneity across the neoplastic and TME compartments – “Senescent Cells,” “Deregulating Cellular Energetics,” “Evading Growth Suppressors,” and “Resisting Cell Death” Cancer Hallmarks. Interestingly, Sibai et al. reported similar ambiguity in these Hallmarks, suggesting that they may reflect patient-specific biological processes and that therapies targeting them may require more individualized treatment selection.

**Figure 2:**
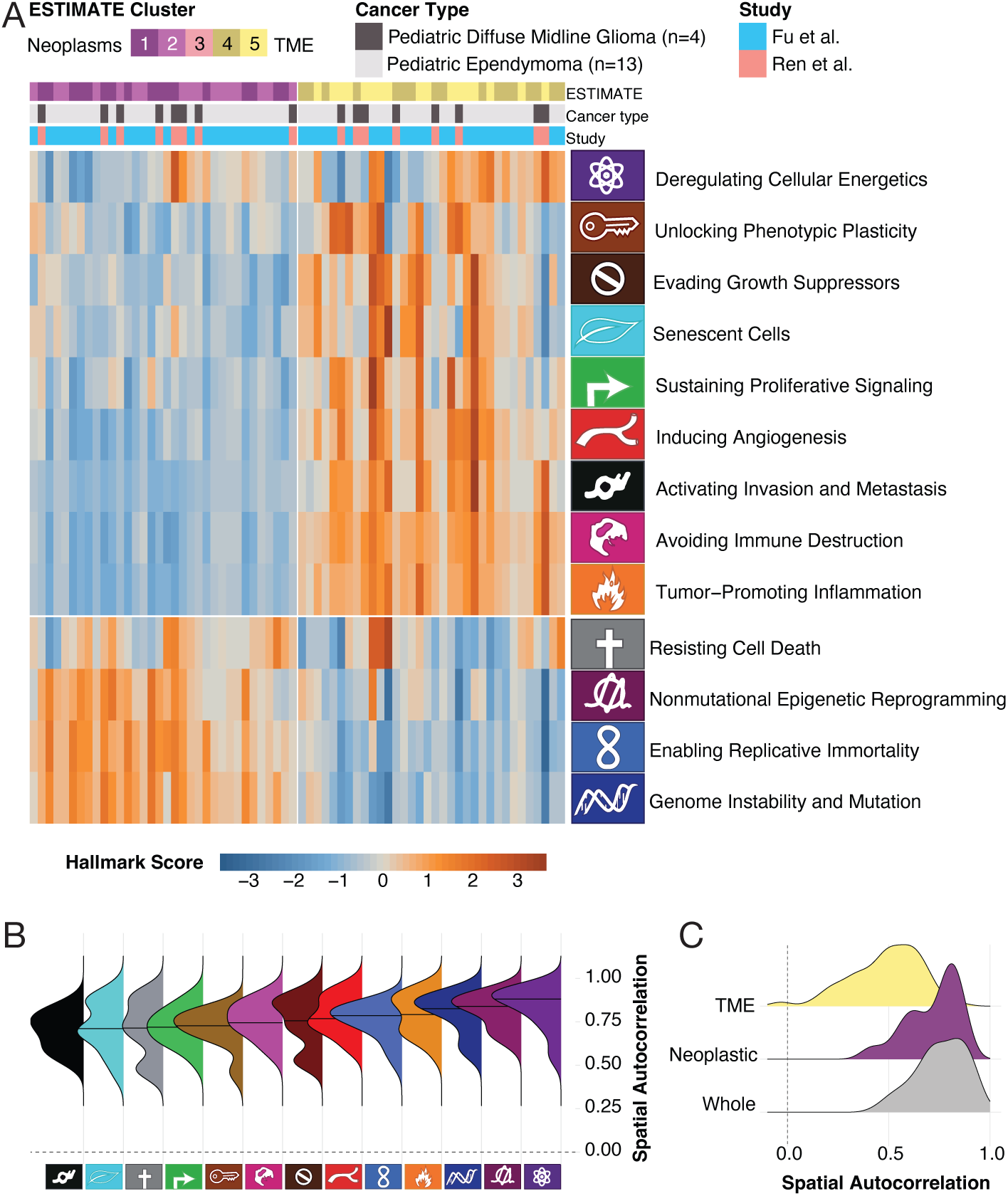
Hallmark Scores by Cancer Neoplastic or TME Compartment. **A)** An unsupervised clustered heatmap of spatial transcriptomics spots by Cancer Hallmark Score (dendrogram removed). Columns indicate sample-compartments as defined by tumor purity (purple is more neoplastic and yellow is more TME). Two additional column annotations include study and pediatric brain cancer type. Rows indicate Cancer Hallmark scores as estimated by gene sets and labels with Hallmark symbols and names. **B)** Distribution of spatial autocorrelation values for each Cancer Hallmark distribution throughout the whole sample is sorted by median from lower spatial organization to higher levels of spatial organization. **C)** Overlapping density plots of spatial autocorrelation of aggregated Hallmarks across each sample. TME compartments, neoplastic compartments, and the whole tissue are displayed.

### SPATIAL AUTOCORRELATION OF GENE EXPRESSION REVEALS HALLMARK STRUCTURE

Spatial autocorrelation analysis, often used in processing geographic data, provides measures that properly account for the non-random nature of neighboring events (Getis 2007). On the molecular level, spatial autocorrelation provides insights into gene expression activity across a tissue section. High positive spatial autocorrelation indicates that neighboring locations exhibit similar biological activity, suggesting the presence of coherent biological regions or cellular neighborhoods. To quantify this pattern, Sibai et al. calculated the spatial autocorrelation of each Cancer Hallmark by correlating its Hallmark score activity vector with a spatial lag vector, which represents the activity of neighboring locations. Values approaching 1 indicate spatial clustering, values near 0 indicate a random spatial distribution, and values approaching −1 indicate that neighboring locations have dissimilar activity. Across adult tumor samples, the average spatial autocorrelation of the 13 Cancer Hallmarks was approximately 0.8, indicating that Hallmark activity was strongly spatially clustered rather than randomly distributed or attributable to a small subset of dominating Hallmark gene sets.

We examined whether this spatial organization was also present in pediatric brain tumors, both across whole samples and within the neoplastic and TME compartments. We observed a similar average spatial autocorrelation across the entire tissue section of pediatric samples (0.76 ± 0.12 SD), suggesting that pediatric brain tumors exhibit broadly similar spatial organization of Cancer Hallmarks. This high degree of spatial organization was observed across all pediatric Cancer Hallmarks (**Fig. 2B**). We next asked whether this spatial organization was maintained within the TME and neoplastic compartments (**Fig. 2C**). Average spatial autocorrelation was lower in the pediatric TME compartment (0.53 ± 0.18 SD) than in the neoplastic compartment (0.68 ± 0.15 SD), a pattern also observed by Sibai et al. in adult tumors.

### GENES DRIVING HALLMARK COMPARTMENT PARTITIONING IN PEDIATRIC AND ADULT BRAIN TUMORS

We examined the gene-level drivers underlying the partitioning of Cancer Hallmark activity between the neoplastic and TME compartments in pediatric versus adult brain tumors (**Materials and Methods**). By comparing fold-expression differences of hallmark-associated genes and contrasting our pediatric data with adult brain tumor profiles from Sibai et al., we identified distinct compartmental shifts in gene expression across age groups (**Fig. 3 and Fig. S3**). We paid particular attention to potential genetic drivers responsible for shifting the three Cancer Hallmarks that no longer clustered with the neoplastic compartment in pediatric brain tumors: “Deregulating Cellular Energetics,” “Senescent Cells,” and “Evading Growth Suppressors”. Several genes emerged as potential drivers of this age-specific compartment switch: PRDM1, TWIST1, GATA2, and IGFBP1 showed strong preference for the pediatric TME (**Fig. 3**) but had no significant compartmental preference in adults glioblastomas (**Fig. S3**). This presence of developmentally regulated transcription factors such as TWIST1, GATA2, and PRDM1 suggests that the pediatric TME may shape unique developmental or immune regulatory cues, reflecting a less differentiated state of stromal and immune cells [PMID: 41446274]. Intriguingly, PRDM1 (aka BLIMP1) has been shown to regulate cell growth by modulating p53 activity (Yan et al. 2007). Furthermore, these genes may also help to explain why immune therapies are often less effective in pediatric tumors (Griesinger et al. 2013; Majzner et al. 2017; Bockmayr et al. 2018; Lin et al. 2018; Lieberman et al. 2019), and suggests that the TME may be less responsive, more immunologically “quiet,” and developmentally protected. Lastly, a large majority of the genes from neoplastic-associated Hallmarks were over-expressed but did not achieve large fold change differences in pediatric nor adult compartments, a result suggesting a tightly regulated network of genes that are subject to active surveillance mechanisms.

**Figure 3:**
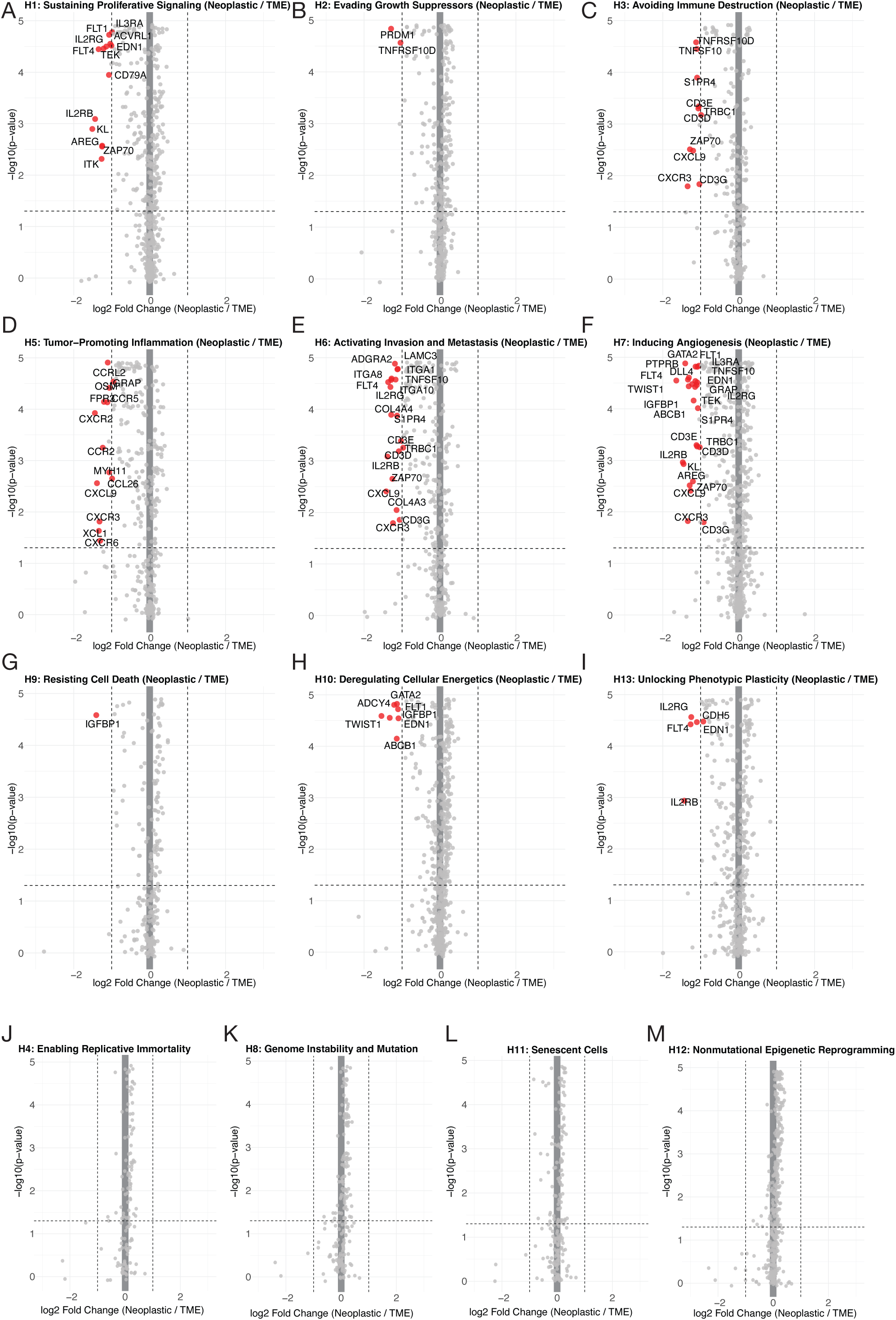
Differential Expression Analysis between Pediatric TME and Neoplastic compartments. Volcano plots comparing mean pediatric Cancer Hallmark scores between neoplastic and TME compartments across samples. Log₂ fold change (neoplastic scores divided by TME scores on the x-axis) is plotted against statistical significance (-log_10_ p-value on the y-axis). Thus, values to the left of zero are more expressed in the TME, and points to the right of zero are more expressed in the neoplastic compartments. Red points indicate significant genes. **A)** Sustaining Proliferative Signaling; **B)** Evading Growth Suppressors; **C)** Avoiding Immune Destruction; **D)** Tumor-Promoting Inflammation; **E)** Activating Invasion and Metastasis; **F)** Inducing Angiogenesis; **G)** Resisting Cell Death; **H)** Deregulating Cellular Energetics; **I)** Unlocking Phenotypic Plasticity; **J)** Enabling Replicative Immortality; **K)** Genome Instability and Mutation; **L)** Senescent Cells; and **M)** Nonmutational Epigenetic Reprogramming.

First, we identified a broad set of genes with >2-fold enrichment in the pediatric TME that showed no such pattern in adults **(Fig. 3 and Fig. S3)**: CCR2, CCR5, CCRL2, CD3D, CD3E, CD3G, CD79A, COL4A3, CXCR2, CXCR3, CXCR6, IL2RG, ITK, KL, S1PR4, TEK, TNFSF10, TRBC1, XCL1, and ZAP70. These genes showed higher expression in the Hallmarks “Sustaining Proliferative Signaling”, “Avoiding Immune Destruction”, “Tumor-Promoting Inflammation”, “Activating Invasion and Metastasis”, “Inducing Angiogenesis”, and “Unlocking Phenotypic Plasticity”. The prominence of chemokine receptors, T- and B-cell signaling components, and vascular-associated genes within this set points to a pediatric TME with coordinated immune cell recruitment, immune signaling, and vascular remodeling that may collectively influence tumor behavior.

We next sought to pinpoint genes that were highly expressed in adult neoplastic compartments yet were also elevated in the pediatric TME—a pattern that could suggest adult tumors must accumulate certain functional traits via mutation that are inherently supported by pediatric TME compartments. Interestingly, the only gene that fit this criterion (with a clear fold expression preference) was CXCL9—a chemokine involved in T-cell recruitment in the body and microglial recruitment in the brain. This observation aligns with emerging models (Vignali et al. 2023; Park et al. 2026) in which exhausted immune cells can form structural and functional barriers that shape tumor progression, such as immunosuppressive stromal barriers that physically or functionally impede effective anti-tumor immunity.

Finally, our analysis of genes driving hallmark partitioning in cancer compartments did not reveal any significantly unregulated genes in neoplastic compartments for any Cancer Hallmarks (**Fig. 3**). This differs considerably from our analysis of adult brain tumors, where several Hallmarks have genes that were seen to be enriched in neoplastic compartments across many hallmarks (**Fig. S3**). This finding adds to the established notion that adult tumors must acquire traits through many drivers (Gröbner et al. 2018) to establish a pro-growth environment and pediatric tumors, with their extremely low mutation rates, do not (Ma et al. 2018).

### ELUCIDATING SPATIAL RELATIONSHIPS BETWEEN TME AND NEOPLASTIC COMPARTMENTS USING RANDOM FOREST FEATURE IMPORTANCE SCORING

Cancer Hallmark phenotypes do not occur in isolation but can interact across the neoplastic and TME compartments (Rinaldi et al. 2018). Rather than functioning as independent programs, the presence and spatial distribution of one phenotype may influence the emergence or maintenance of other phenotypes in neighboring compartments. For example, genetic distance has been shown to correlate with spatial distance in brain gliomas (Gates et al. 2019). As such, spatial relationships may be particularly relevant to how tumor and TME programs are organized. We thus sought to characterize the spatial interdependence of Cancer Hallmarks across compartment boundaries, asking whether the spatial distribution of one Hallmark could predict the distribution of another. Specifically, we aimed to identify which Hallmarks in the neoplastic or TME compartment were most informative of Hallmark activity in the opposing compartment, thereby highlighting potential cross-compartment signals that may contribute to the organization of tumor-associated phenotypes. To make the multifarious components of this section clearer, we divided it into subsections with additional subheadings.

### SCORING COMPARTMENTS TO FACILITATE CROSS COMPARTMENT COMPARISONS

Before we could model spatial interactions between compartments, we interpolated TME hallmark scores onto Neoplastic spots, and Neoplastic hallmark scores onto TME spots. For every subspot of one compartment, we computed an inverse-distance weighted sum of hallmark scores from the opposing compartment and called the new value a “radar score” **(Fig. 4A)**. This process was repeated for all neoplastic subspots to model the spatial impact of TME Cancer Hallmark activity on the neoplastic compartment visa versa. Thus, with radar scores from each Hallmark projected on the opposing compartment we could then estimate the relative importance of each hallmark on the neighboring Hallmark as a way to measure interactions.

**Figure 4:**
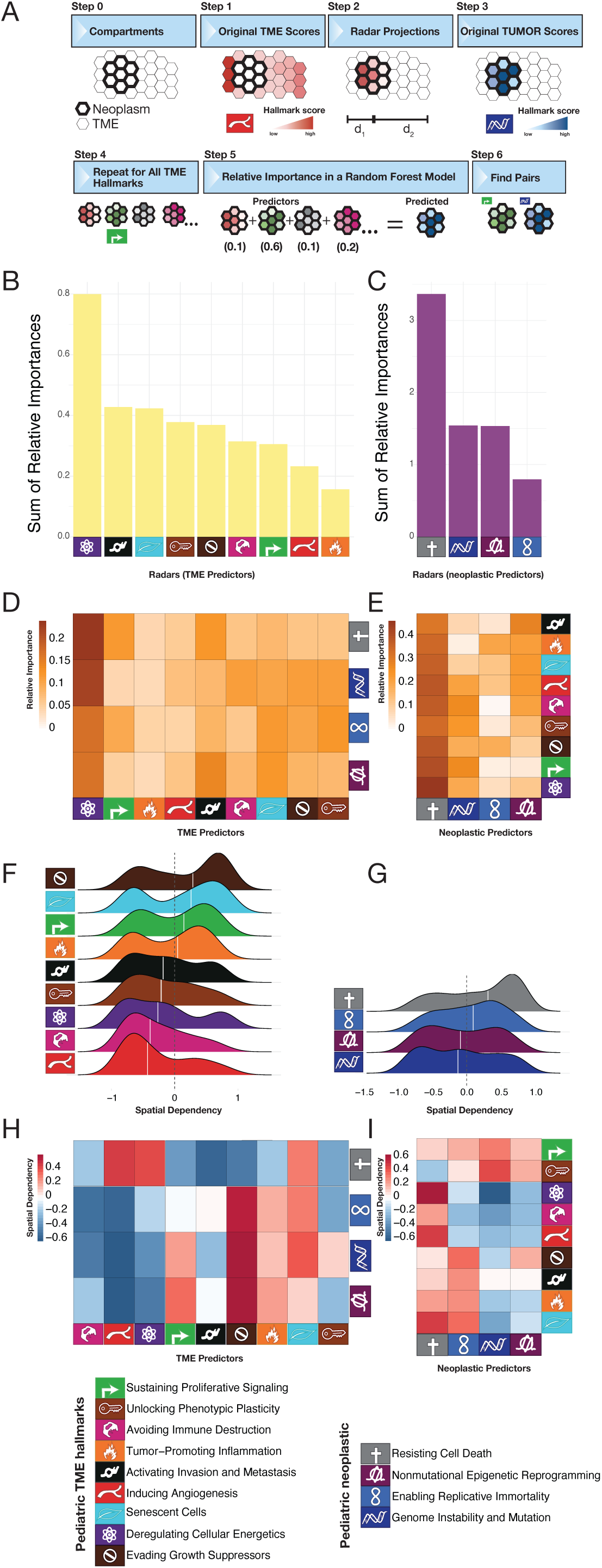
Spatial Dependencies and Model-Predictor Importances Across Tumor Compartments. **A)** Analytical workflow to establish compartment-specific hallmark scoring for cross compartment modeling. <u>Step 0:</u> An illustration shows sample transcriptomic subspots divided into two compartments, neoplastic and TME. <u>Step 1:</u> Hallmark expression scores are highlighted in one compartment. <u>Step 2:</u> A visual representation of projecting one compartment onto another. d1 and d2 illustrate that each compartment projection is weighted by the distance between subspots. These projections will be used as predictors in our model. <u>Step 3:</u> Shows that the targets of the predictors are the original hallmark scores. <u>Step 4:</u> Illustrates a repeat of step 2 for each hallmark to populate the features of the model. <u>Step 5:</u> Illustrates how each hallmark from the opposing compartment was included in the model to predict one Hallmark. <u>Step 6:</u> Evaluate pairs of the hallmarks from the model for possible cross compartment Hallmark interactions. **B)** A bar chart shows the summed relative importance of TME radar predictors, i.e., features of the model. **C)** A bar chart shows the summed relative importance of neoplastic radar predictors. **D)** A heatmap shows TME predictor-importance across neoplastic hallmarks. Predictor-Hallmarks are on the x-axis, and the predicted-Hallmarks are on the y-axis. **E)** A heatmap indicates Neoplastic predictor-importance heatmap across TME hallmarks. Again, Predictor-Hallmarks are on the x-axis, and the predicted-Hallmarks are on the y-axis. **F)** A ridgeplot displays spatial dependency distributions for TME hallmarks. **G)** A ridgeplot displays spatial dependency distributions for neoplastic hallmarks. **H)** A heatmap shows Pearson correlation across subspots between the model’s radar score and its SHAP value in the target-hallmark model. Red values indicate Hallmarks that tend to be active near one another. Blue values tend to be distant from one another. TME predictors are on the x-axis, and the predicted neoplastic relationships are on the y-axis. **I)** Same as panel H but with swapped TME and neoplastic axes.

### RANDOM FOREST MODELING TO IDENTIFY THE CROSS-COMPARTMENT INTERDEPENDENCE OF CANCER HALLMARKS

To determine which Cancer Hallmarks most strongly correlate spatially with each other across compartment transitions, we used impurity-based random forest models to assess how well projected radar scores can predict Hallmark activity in the other compartment. For each sample, we trained a model on 80% of the target subspots’ radar scores and hallmark scores, and then evaluated its performance on the remaining 20% of subspots. We found our models to be effective at explaining the variability of cancer hallmark scores (mean R² of 0.69±0.12 standard deviation across all samples). Our mean R² value was lower than that observed by Sibai et al. (0.82).

### IDENTIFYING RELATIVE IMPORTANCE OF CANCER HALLMARKS BY PREDICTING HALLMARK SCORES FOR EACH OPPOSING COMPARTMENT HALLMARK

Because impurity-based random forest models can bias features with more variability (Strobl et al. 2007), we used Shapley additive explanations (SHAP) – which provides less biased feature attribution – to better understand our random forest models. SHAP values quantify how much each feature contributes to an individual prediction relative to the model’s baseline prediction. We used SHAP values to analyze relative importance and spatial dependency of hallmark interactions (**Samples and Methods**). Relative importance assesses which predictor hallmarks are the biggest drivers of hallmark activity in the other compartment, whereas spatial dependency assesses whether these interactions are driven by spatial proximity.

Examining the relative importance of each predictor Cancer Hallmark in each model, we observed that “Resisting Cell Death” had highest relative importance among radars originating from the neoplastic compartment, and that “Deregulating Cellular Energetics” and “Activating Invasion and Metastasis” had the highest relative importances among radars originating from the TME compartment (**Fig. 4B** and **Fig. 4C**). Of note, both “Resisting Cell Death” and Deregulating Cellular Energetics” had some of the cross compartment scores (**Fig. 2A**). When we ran the model without some of these cross compartment Hallmarks, we observed strong model contributions from the “Inducing Angiogenesis” and “Tumor−Promoting Inflammation” (**Fig S4**) Seeking to determine individual relationships between Cancer Hallmarks, we further investigated the relative importances between each TME Cancer Hallmark and each neoplastic Cancer Hallmark (**Fig. 4D** and **Fig. 4E**).

When examining the relative importance of neoplastic Cancer Hallmarks affecting TME Cancer Hallmarks, we observed strong relative importance scores from the “Resisting Cell Death” gene sets across all TME Hallmarks (**Fig. 4E**). We also observed that “Nonmutational Epigenetic Reprogramming” exerted strong effects on every TME cancer hallmark except for “Sustaining Proliferative Signaling.” In contrast, “Enabling Replicative Immortality” had minimal importance in predicting any of the TME Cancer Hallmarks (**Fig 4E)**. Additionally, “Genome Instability and Mutation” exerted strong effects on both “Sustaining Proliferative Signaling” and “Avoiding Immune Destruction.” (**Fig. 4D**). Again when restricting this analysis to fewer Hallmarks, “Enabling Replicative Immortality” remained the weakest predictor of TME (**Fig. S4C and S4E**).

When examining the relative importance of TME Cancer Hallmarks affecting the neoplastic Cancer Hallmarks, we observed that only three hallmarks were meaningful in predicting the tumor’s hallmarks. “Deregulating Cellular Energetics” was a strong predictor of “Resisting Cell Death” and “Genomic Instability”. Whereas “Activating Invasion and Metastasis” were influential in predicting “Nonmutational Epigenetic Reprogramming” and “Resisting Cell Death” (**Fig 4B and 4D**). Interestingly, both of these neoplastic Cancer Hallmarks were not affected by “Tumor-Promoting Inflammation” when all Hallmarks were included in the model but showed stronger effects when we excluded the top predictors, suggesting intracompartmental interactions (**Fig S4B and S4D**).

### EVALUATING THE SPATIAL DEPENDENCY OF CROSS-COMPARTMENT INTERACTIONS

We next asked whether the influence of Cancer Hallmark predictors was associated with their spatial context. For each target Cancer Hallmark and each predictor Hallmark, we calculated a spatial dependency score as the Pearson correlation across subspots between the predictor’s radar score and its SHAP value in the target-hallmark model (**Samples and Methods**). Thus, this metric, because of the nature of the distance weight of the radar-scores, quantifies whether variation in the spatial relationship contributes to target-hallmark prediction. Unlike spatial autocorrelation, which measures whether a variable exhibits spatial clustering or similarity among neighboring locations, our spatial dependency score specifically measures whether a predictor’s spatial context is associated with its influence on the prediction of another Hallmark. Positive spatial dependency means the predictor and target hallmarks tend to be active near one another along their shared interface. Negative spatial dependency means they tend to occur farther apart. Nonlinear spatial dependencies may reflect more localized patterns, with the hallmarks aligned in particular regions near to or distant from the interface. After calculating the spatial dependencies for each hallmark combination, we took the median spatial dependencies across all samples (**Fig. 4F-4I, Fig S4F-S4I**).

Many Cancer Hallmarks had bi-modal distributions of spatial dependence (**Fig 4F-4G and Fig S4F-S4G**) suggesting strong proximal and distal impacts in cross compartment relationship. When looking at inter-compartment comparison of Hallmarks, we observed “Evading Growth Suppressors” to have a strong spatial relationship with most Neoplastic Hallmarks (**Fig. 4H**) We also observed that “Inducing Angiogenesis” had negative spatial dependencies for every neoplastic Cancer Hallmark (**Fig. 4H**). Rather than indicating that angiogenesis is unrelated to pediatric tumor progression, these negative associations may reflect spatial specialization of the tumor microenvironment, in which immune-evasive and vascular-associated programs are concentrated in niches that differ from those supporting the strongest tumor-intrinsic Cancer Hallmark programs. When we removed the two top predictors, we noticed that the spatial dependencies of our pediatric tumor samples contained many similarities to Sibai et al.’s adult tumor samples. For example, when we examined the spatial dependencies of signals originating from the TME compartment, we observed that “Sustaining Proliferative Signaling” had highly positive spatial dependencies for every neoplastic Cancer Hallmark (**Fig. S4**). This consistent positive association suggests that proliferative signaling may identify tumor niches in which multiple tumor-intrinsic programs are simultaneously reinforced. Such spatial coupling is consistent with emerging evidence that glioma tumor states are organized into distinct microanatomical niches rather than being uniformly distributed throughout the tumor(Greenwald et al. 2024).

When we examined the spatial dependencies of signals originating from the neoplastic compartment, we observed that each one shared a strong positive spatial dependency on “Sustaining Proliferative Signaling,” a pattern that held when the model included all hallmarks and that restricted model (**Fig 4I and Fig. S4I, respectively**). We observed that “Genome Instability and Mutation” within the neoplastic compartment is largely inhibitory of Cancer Hallmarks in the TME and is also inhibited by many Cancer Hallmarks in the TME. Likewise, similar to the adult samples, we also observed that the main neoplastic driver of cancer hallmark behavior in the pediatric TME appears to be “Nonmutational Epigenetic Reprogramming.” Whereas neoplastic “Nonmutational Epigenetic Reprogramming” promotes TME “Sustaining Proliferative Signaling” in both pediatric and adult samples, it also promotes pediatric TME “Unlocking Phenotypic Plasticity.” (**Fig 4I and Fig. S4I)** This difference is unsurprising since children are already in a developmental state involving cellular differentiation (Lawlor and Thiele 2012). We also noticed “Avoiding Immune Destruction” was inhibitory of neoplastic Cancer Hallmarks. We found this finding interesting because pediatric brain tumors are often considered immunologically cold (Gröbner et al. 2018).

### INTER-TUMOR AND INTRA-TUMOR SPATIAL DEPENDENCE AND INDEPENDENCE

Adding to the previous section, we wanted to identify strong pair-wise examples of Hallmark interactions and gain greater insights into which hallmarks are most spatially dependent and independent. Using the spatial dependency scores, as described in the previous sections, we aggregated spatial dependency scores and found, when attempting to predict Hallmark expression in the TME (**Fig. 5A**) and vice versa (**Fig.5B**), that neoplastic hallmarks have greater proximal effect on the TME than the TME on the tumor. To illustrate these pairwise comparisons we used a co-correlation graph where spot-level Hallmark scores are plotted. When the scaled expression scores are both high, then the plots are colored white. When spots show low scores in both Hallmarks colors go black. If the expression score of one Hallmark is greater than another, blue or orange colors are shown (**Fig. 5C** and **Fig. 5D**). Using one ependymoma sample, we highlight some of the most extreme spatially dependent pairwise Hallmarks.

**Figure 5:**
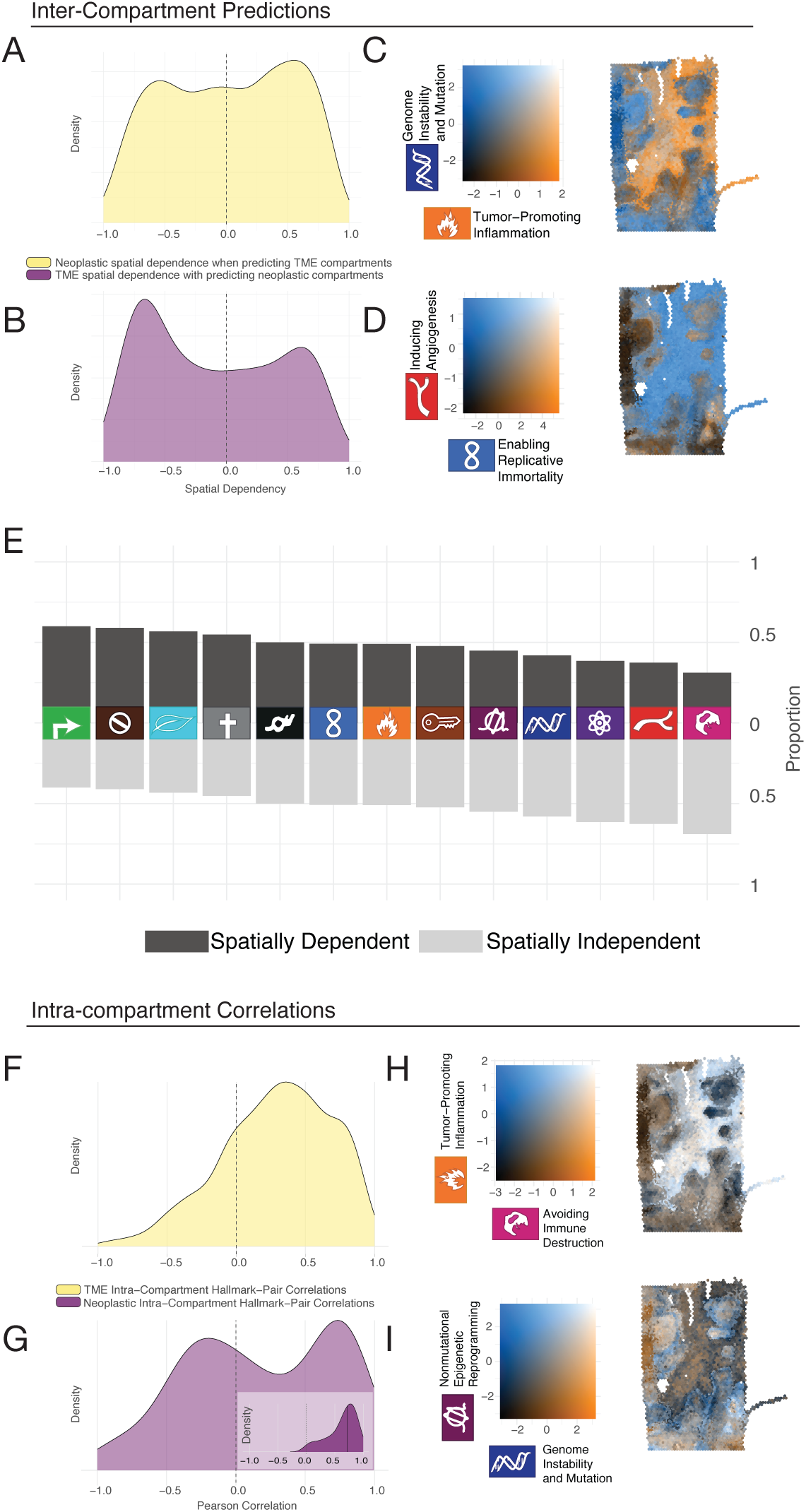
Examples of spatial dependency for intra- and inter-compartment hallmark relationships. **A)** A density plot illustrates the distribution of spatial dependency sample-level for each pair-wise hallmark. The top density plot (yellow) displays the per-sample median spatial dependency scores for all pairs of neoplastic predictors that predict a spatial relationship with TME hallmark scores. High scores represent strong spatial dependence (similar expression in close proximity), whereas lower values indicate distal correlations. **B)** Similar to Panel A, this density plot (purple) displays the per-sample median spatial dependency scores for all pairs of TME predictors that correlate with TME hallmark scores. **C)** Example of colocalized scaled hallmark expression plots between intercompartmental hallmarks “Genomic Instability” and “Tumor-promoting inflammation”. In the color legend, colocalized hallmarks at maximum activity are shown in white, and colocalized minimal activity in black. Orange and blue indicate the activeness of one hallmark with little to no expression of the other. Colocalized hallmark activities appear in white and black, while orange and blue shows compartment specific findings. **D)** Same color scheme as Panel C for two different hallmarks **E)** A likert-style plot highlights the proportion of Hallmarks with strong spatial dependence scores (>0.2) as predictors and targets suggesting which Cancer Hallmarks have strong spatial dependence. **F)** Density plot of intra-compartment paired correlations scores using intro compartment correlations estimates in the TME. **G)** Same as Panel F, but for Neoplastic compartments. **H & I)** Same as Panels C and D but for intra-compartment correlation scores.

Interestingly, when we counted the number of pair-wise interactions with greater than 0.2 (correlation coefficients) as a metric of spatially proximal co-regulations, we observed that “Proliferative signaling”, followed by “Evading Growth Suppressors”, were the Hallmarks most spatially dependent on their neighbors (**Fig. 5E**). Interestingly, “Angiogenesis” and “Avoiding Immune Destruction” tended to have the lowest dependency scores overall. Whether this is due to existing vasculature the remains unaffected or the mobility of immune cells warrant further investigation.

To evaluate Cancer Hallmark correlation within neoplastic and TME compartments, we calculated pairwise Pearson correlations for subspots in each compartment across all samples. We calculated correlation scores for each set of pairs in the neoplastic and TME hallmarks, then summarized the correlations within each sample by their mean and mean absolute value. We visualized the distributions of pairwise correlations across samples using density plots (**Fig. 5F and 5G**). To assess how the neoplastic correlation distribution changed when individual hallmarks were excluded, we generated additional plots omitting, in turn, all pairs that included each neoplastic hallmark (**Fig. 5G-insert**). This removal of one Hallmark, “Resisting Cell Death”, resulted in strong coregulation of the remaining Neoplastic Hallmarks, “Nonmutational Epigenetic Reprogramming”, “Genome Instability and Mutations”, and “Enabling Replication Immortality”, which are closely and tightly coregulated within Neoplastic Compartments.

To illustrate these pairwise relationships within microenvironmental (**Fig. 5H**) and neoplastic (**Fig. 5I**) compartments, we looked at one ependymoma sample using the co-correlation plots, the same sample as above. As illustrated by the abundance of black and white coloration when observing the relationship between “Tumor-Promoting Inflammation” and “Avoiding Immune Destruction,” this indicates these two Hallmarks are co-regulated in the TME. Interestingly, when we looked at the co-correlation of “Nonmutational Epigenetic Reprogramming” and “Genome Instability and Mutations”, two signature-Hallmarks of Neoplasm compartments, we observed a unique pattern where the fringes or borders of the tumor regions expressed genes specific to “Nonmutational Epigenetic Reprogramming” Hallmark, whereas “Genome Instability and Mutations” Hallmarks genes occupied the deeper tumor.

### NO SUBCLONES WERE IDENTIFIED

Sibai et al. used inferCNV (Tickle et al. 2019) to group subspots into likely clonal groups according to copy number profiles. They found that genetic distance between adult tumor clones was correlated with larger differences on average in neoplastic Cancer Hallmark activity. However, when we performed this same analysis on our pediatric data, we did not observe any CNV-governed subclones. This result likely reflects a genomically stable and hypomutated state of our pediatric brain tumors. However, the sequencing resolution of 10X Visium is such that we cannot exclude extensive polyclonal diversification as was previously observed in pediatric Wilms tumors using single molecule mutation detection from nanoseq duplex sequencing (Lee-Six et al. 2025).

### ASSOCIATION OF TUMOR GRADE AND HALLMARKS ON TUMOR PURITY

Although tumor purity minimally affects overall survival among pediatric CNS tumor patients, (Yang et al. 2021), we hypothesized that some genes may be strongly associated with the auto correlation of tumor purity scores and provide clear rationale for using spatial transcriptomics more commonly for tumor pathology.

Here, we quantified a sample-level tumor purity score by calculating the spatial autocorrelation of ESTIMATE scores across subspots (as reported earlier) (Aran et al. 2015). This value captures how consistently subspots with similar ESTIMATE scores are located near one another, with higher values indicating greater spatial organization of regions with similar tumor and microenvironmental composition. We calculated this measure for all subspots combined and used the resulting spatial autocorrelation as our sample-level measure of tumor purity and called it a Spatial Coherence Score (SCS).

Very few clinical variables were provided for these published CNS tumors to evaluate associations with our SCSs (**Supplementary Table 1**). Tumor grade, however, was available for a subset of our tumors (the ependymoma samples from Fu et al., 2023). When we examined the relationship between a sample’s tumor WHO grade (Torp et al. 2022) and its SCS, we observed a clear association suggesting that SCS may have clinical utility as a digital pathology measure (**Fig. 6A**, p-value=7.0×10^−3^, Student’s t-test). Here we interpret that higher SCS to be more organized and differentiated, aligning with the definition of tumor grade.

**Figure 6:**
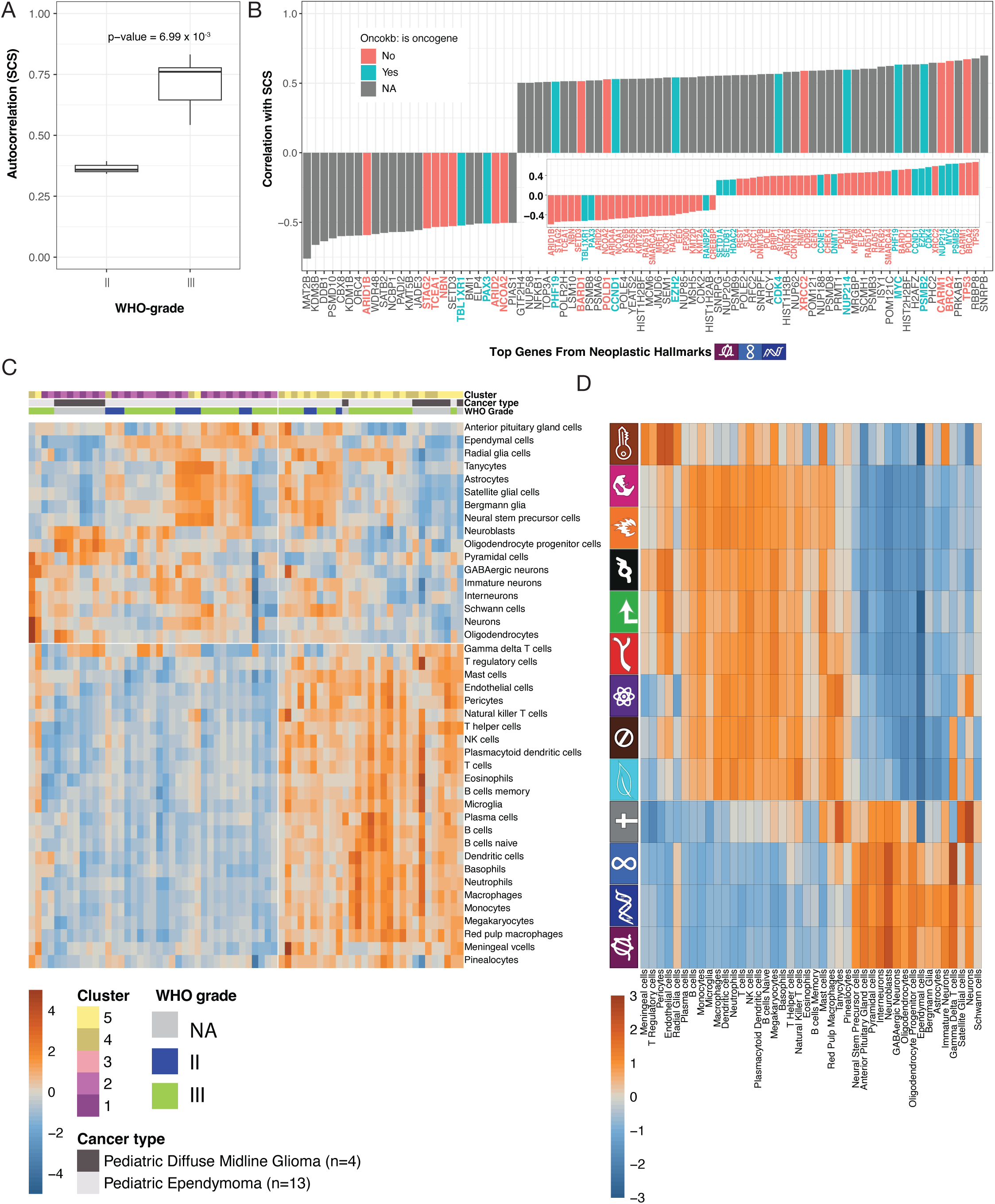
Correlation of Spatial Coherence Score (SCS) with tumor grade and gene-level expression from Neoplastic Hallmarks. **A)** A Boxplot indicates the Spatial Coherence Scores (SCS or the Spatial Autocorrelation of ESTIMATE scores indicating tumor purity) on the y-axis and ependymoma WHO-tumor grade on the x-axis. **B)** A barplot of the top correlations, i.e., greater than 0.50 or less than −0.50 between SCS and genes from Neoplastic Hallmark genesets. Bars are colored by oncoKB annotation of tumor suppressor (orange) or oncogene (blue). The inset plot is restricted to oncoKB annotated genes at a new correlation cutoff > 0.3 or a cutoff < −0.3. **C)** Heatmap showing cell-type scores across neoplastic and TME compartments. Cell-type scores were calculated within each sample, averaged across all samples, and scaled across sample-compartment observations for visualization. Study and cancer type are also annotated. **D)** Heatmap showing cell-type scores across Cancer Hallmarks. Cell-type scores were calculated within each sample, averaged across all samples, and scaled across each Hallmark for visualization.

When applied to our Cancer Hallmark Framework, we observed that no single Cancer Hallmark was clearly correlated with SCS, and by extension the tumor’s grade (data not shown). As such, we investigated correlations at the individual gene level within our neoplastic-associated hallmark gene sets (**Fig. 6B**), asking which Neoplastic Hallmark-associated genes correlated best with the samples’ overall SCS or grade. Limiting to genes with Pearson correlations greater than 0.5 or less than −0.5 from the “Genomic Instability”, “Nonmutational Epigenetic Reprogramming”, and “Enabling Replicative Immortality” (a total of 590 genes), we identified many candidate genes that tend to associate with SCS (**Fig 6B**). When annotated with the oncoKB cancer gene list (n=139), we observed TP53, MYC, PSMB2, and CARM1 expression to be strongly associated with SCS and by extension higher tumor grades. Conversely, lower expression of ARID1B and others were correlated with higher tumor grade. As expected, we see overexpressed genes to have more oncogenes (**Fig 6B inset**) and more tumor suppressor genes to be negatively associated with SCS, as annotated by OncoKB (Chakravarty et al. 2017).

### DECONVOLUTION OF THE CANCER HALLMARKS INTO PREDICTED CELL TYPES REVEALS CANCER TYPE SPECIFIC ORIGINS AND TME COMPOSITION

At the most fundamental level, hallmark-level patterns suggest that the composition of each compartment reflects the contributions of its constituent cell types. To evaluate cell type contributions to compartments and Hallmark-level observations we used scRNA marker gene signatures, curated by PanglaoDB (Franzén et al., 2019), to evaluate cell composition (**See Methods**). Unsupervised clustering of the resulting cell type scores revealed a distinct separation between the TME and neoplastic compartments (**Supp. Fig. 6C**). The correspondence between the inferred cell type clusters and the previously established compartment assignments was statistically significant (Fisher’s exact test, p = 4.047 × 10⁻¹⁴), supporting the ability of these cell type signatures to distinguish the two spatial compartments. Examination of the cell type distribution revealed a striking compartment-specific organization. Immune and vascular cell types preferentially clustered within the TME, consistent with the expected enrichment of immune and stromal populations within the tumor microenvironment. In contrast, the majority of the 20 nervous system cell types examined preferentially associated with the neoplastic compartment, suggesting that the neoplastic compartment retains a transcriptional identity enriched for neural cell type signatures. Upon closer inspection of the neoplastic compartment, we observed that tumor-type also clustered together highlighting distinct tumor cell of origin. Specifically, neuroblasts and oligodendrocyte progenitor cells expression signatures were more present in the diffuse midline gliomas study. In the 26 Neoplastic compartments from the ependymoma study, we observed the strongest cellular expression signatures in ependymal cells and radial glia cells (**Fig 6C**).

Finally, we sought to explore the relationship of how cell-type composition correlated with Cancer Hallmark activity (**Fig 6D**). Our analysis showed that the activity of “Sustaining Proliferative Signaling”, “Avoiding Immune Destruction”, “Tumor-Promoting Inflammation”, “Activating Invasion and Metastasis”, and “Inducing Angiogenesis” strongly associated with immune cells, microglia cells, and vasculature cells (endothelial cells, pericytes). We also observed neoplastic strong activity of “Enabling Replicative Immortality”, “Genome Instability and Mutation”, and “Nonmutational Epigenetic Reprogramming” associated with nervous system cell types. Immune cells, tanycytes, pinealocytes, and neurons were found to be active in “Evading Growth Suppressors”, “Deregulating Cellular Energetics”, and “Senescent Cells” Hallmarks. “Resisting Cell Death” correlated with nervous system cell types, and several immune cell types (red pulp macrophages, mast cells, T helper cells, neutrophils, basophils, natural killer T cells, megakaryocytes, B cells naive, NK cells); and “Unlocking Phenotypic Plasticity” also correlated with immune cells, but also bergmann glia cells, immature neurons, and schwann cells (**Fig 6D**).

## Discussion

First described by Hanahan and Weinberg in 2001 (D. Hanahan and Weinberg 2000; Douglas Hanahan and Weinberg 2011; Douglas Hanahan 2022), the Cancer Hallmarks offer a high-level framework for organizing the complex cellular interactions that drive and sustain tumor progression. Building on the work of Sabi et al., which used a Cancer Hallmark framework to examine spatial transcriptomics across 63 adult tumors representing 10 cancer types (Sibai et al. 2025), we now investigate pediatric brain tumors. Given their low mutation burden (Rahal et al. 2018) and developmentally active (or stalled) milieu (Gonzalez Castro et al. 2023; Jessa et al. 2019), we asked how pediatric brain tumors differ in Cancer Hallmark activity across the neoplastic and tumor microenvironment (TME) compartments. Although adult and pediatric brain tumors share many features, we repeatedly observed greater hallmark activity in the TME of pediatric samples when compared with the TME of adult samples. We also found that simple image-analysis metrics of spatial autocorrelation can be applied to transcriptomic data to assess tumor WHO grade.

Spatial transcriptomics is rapidly becoming a powerful tool that enables researchers to examine the impact of ecological communities in tissue samples (Ståhl et al. 2016; Lewis et al. 2021). Here, Hanahan and Weinberg’s Cancer Hallmarks provide a high-level interpretation of tumor and neoplastic interactions. While this approach is less valuable in determining single-gene biomarkers, it does provide an intriguing evaluation of the TME’s contribution to sustaining tumor growth. Here we show, when compared to the adult glioblastoma, that pediatric tumors do not have nearly as many cancer Hallmarks represented in Neoplastic compartments but are far more expressive in TME compartments. Specifically, we observed changes in “Evading Growth Suppressors”, “Deregulating Cellular Energetics”, and “Senescent Cells” Hallmarks which are more expressed in Adult Neoplastic compartments. These differences may point to key functions that must be acquired over time through somatic mutations (Lawrence et al. 2013) to generate, shape, or mimic the extracellular signaling naturally present in the developing brains of pediatric patients.

While pediatric tumors expressed Hallmarks very differently from their adult counterparts, many of the genes that were “overexpressed” in Neoplastic associated Hallmarks were not significantly different than their TME counterparts. This non-significant overexpression was observed in both pediatric and adult cohorts (**Fig. 3** and **Fig. S3**, respectively). We find this result to potentially reflect the delicate evolutionary constraints of cancer growth in a system with many checkpoints for programmed cell death and immune regulation. Of the ∼600 genes associated with the three main neoplastic Hallmarks: “Non-epigenetic Reprogramming”, “Genome Instability and Mutation”, and “Enabling Replicative Immortality” only 4 genes in adults showed significant fold change differences (>2X) between neoplastic and TME compartments. Interestingly, in nearly every gene connected to neoplastic Hallmarks we observed overexpression relative to TME compartments; this is true in both the pediatric (**Fig. 3)** and adult samples (**Fig. S3**). We interpret this striking observation with many possible hypotheses. First, this collective increase of expression across many genes remains undetectable to governors of cell growth because evolutionary mechanisms work on a gene-by-gene and protein-by-protein interaction level and no master regulator has evolved for polygenic signals from hundreds of slightly upregulated genes. Another hypothesis follows the rationale for the molecular interpretation for the relative success of platinum agents and chemotherapy in treating tumors globally. By pushing cells into higher expression states of DNA damage, replicative immorality, and epigenetic reprogramming, cellular governors of growth are activated and result in effective treatments. Each of these hypotheses will require greater attention and experimental validation, but point to a cancer’s ability to skirt the boundaries of gene expression to remain undetected and achieve success in replication by creating cancer with a community of cells.

One of the foundational and potentially never-resolved questions in cancer biology is how cells react to their neighbors and environments. Our distance-based radar scores analysis was one of many possible approaches to address this relationship and the interactions between TME and neoplastic environments. Some striking examples showed strong intra-compartment expression similarities with “Genome Instability and Mutation” Hallmark at a tumor compartment’s core compared to “Nonmutational Epigenetic Reprogramming” which showed more readily at the tumor boundaries. And coregulation from the inter-comparison also shows very strong associations between “Tumor-Promoting Inflammation” and “Avoiding Immune Destruction” with the inflammation-Hallmark in closer proximity to the tumor boundary. We believe that with improved experimental design, incorporating time-course data we will start to explore the direction of effect of these interactions to better understand tumorigenesis and the order of operations that lead to the earliest signs of cancer.

## Limitations

While our study offers a unique examination of the behaviors of pediatric brain cancers, it is limited by several factors. While there exist a multitude of pediatric brain cancers, spatial transcriptomic data from pediatric brain cancer patients was limited at the start of this study. As such, our analysis only included two forms of pediatric brain cancer, and drew on a modest sample size. We anticipate that as spatial transcriptomic data become more available in coming years, our understanding of pediatric brain cancer ecology will grow. Another limiting factor of our study was the inherent lack of single-cell resolution provided by VISIUM. We attempted to abate these concerns by approximating single-cell resolution with BayesSpace. Another potential limiting factor of our study was the use of different versions of Space Ranger (v1.1.0 and v2.1.1), although we don’t believe that this discrepancy significantly altered our study findings. We also recognize that several confounding factors likely influenced our analysis. For instance, much of our analysis relied extensively on radar scores and random forest modeling. However, the accuracy of radar scores fluctuates with several factors, such as tumor purity, tumor size, and tumor position on the VISIUM slide. With such variability, our study is best understood as identifying high-level, general trends seen in pediatric brain tumors. Of note, mutations within the sample in Visium v2 assays can potentially interfere with appropriate probe hybridization. Specifically, mutations that occur within the 50 base pair gene-specific target sequence (25 bp in the Left-Hand Side Probe, 25 bp in the Right-Hand Side Probe) could negatively impact transcript detection, although some variation is tolerated. Mutations that cause a mismatch at the ligation junction between probes cause a decrease in ligation efficiency. (10xgenomics.com) Thus, in highly mutated tumor samples, target region modifications may be a non-trivial concern with data efficacy. However, given the lower mutation burden in pediatric tumors we have assumed high-efficacy gene identification in Visium experiments.

This study does not include a comprehensive collection of pediatric brain tumors. Yet, of the many central nervous system tumors that occur (Kline and Sevier 2003)), it is estimated 9% are ependymomas(Gurney 1999) and 10-15% are diffuse midline gliomas(Nikbakht et al. 2016). Collectively these two tumor types account for more than one-fifth of new pediatric brain tumors in the United States. The median patient age at ependymoma diagnosis is five years old, with 25-40% of ependymoma patients younger than two years old (Formentin et al. 2023). This age distribution is seen in many forms of pediatric brain cancer, like medulloblastomas. Thus, the pediatric ependymoma model is informative of the impact that key developmental processes have on tumorigenesis. Regarding location, pediatric ependymomas most commonly occur in the posterior fossa, which hosts 45-60% of pediatric brain lesions (Formentin et al. 2023). The most common brain tumors of the posterior fossa are ependymoma, medulloblastoma, and pilocytic astrocytomas. Therefore, the pediatric ependymoma model shares similar microenvironments and interactions with surrounding brain tissue as many other pediatric brain tumors (Formentin et al. 2023).

Cancer type is an inevitable confounding factor that we expected would influence several Cancer Hallmarks. In particular, we anticipated differences in the “Activating Invasion and Metastasis” and “Inducing Angiogenesis” Cancer Hallmarks. While pediatric ependymoma originates from glial cells (pediatric ependymoma originates in ependymal cells in the ventricular system), it forms in different regions of the brain. Pediatric ependymomas form primarily in the cerebellum and spinal cord. In adult brains, astrocytes are very plentiful and tile the entire CNS, while ependymal cells are absent from large swaths of the lateral ventricles (Del Bigio 2010). Pediatric ependymoma is typically less invasive and forms well-defined tumor masses. Because pediatric ependymoma is less vascularized, we expected that it would have less expression of the “Inducing Angiogenesis” Cancer Hallmark. Despite anticipated differences between diffuse midline gliomas and ependymomas, we saw very little effect at the Hallmark-level.

## Declaration of Generative AI in the Research and Writing Process

During the research process, the writers used ChatGPT to debug code and write several code scripts. The authors reviewed the code before and after use and take full responsibility for the code used in the analysis. In the writing process, the authors also used an LLM to articulate hypotheses and Perplexity.ai for literature searches, followed up by human fact-checking within published literature found on Google Scholar. The authors also used LLMs like ChatGPT to revise manuscript excerpts for clarity and facilitate the conversion of ideas into language. The authors reviewed and edited the suggestions and take full responsibility for the content of this publication.

## Acknowledgements

This project was supported by funding from Brigham Young University’s Simmons Center for Cancer Research Interdisciplinary Research Award. The authors are grateful to the patients whose samples made this study possible, as well as Fu et al. and Ren et al. for providing publicly available spatial transcriptomic data. The authors also thank Brigham Young University’s Office of Research Computing for providing the high-performance computing resources necessary for the analyses.

## Author Contributions

M.S., M.H.B., E.P-P., S.C., D.G., and N.T.M. contributed to conceptualization. M.S., M.H.B., S.C., and D.G. contributed to methodology. C.T.T. was responsible for data curation. N.T.M. contributed to software development. M.H.B., N.T.M., C.W.M., and C.T.T. performed the formal analysis. M.H.B., N.T.M., and C.T.T. contributed to visualization. All authors contributed to writing and revising the manuscript.

## Materials and Methods

The majority of our analysis is modeled after Sibai et al. Here we give a brief overview of the methods used and emphasize where our analysis differed from Sibai et al. For a more detailed explanation of the methods employed, the reader is referred to Sibai et al. and our code repository.

### Data

Using publicly available repositories, we obtained spatial transcriptomic data from 17 pediatric brain cancer samples representing two tumor types: ependymoma (*n* = 13) and diffuse midline glioma (DMG; *n* = 4). The ependymoma data were generated by Fu et al. (2023) and deposited in the NCBI Gene Expression Omnibus (GEO) under accession number GSE195661. Although the original study included 14 pediatric ependymoma samples, one sample was excluded from our analysis due to extreme compartment hallmark expression (**Fig. S2A**), leaving 13 samples for downstream analyses. The DMG data generated by Ren et al. were obtained from the NCBI GEO under accession number GSE194329. Some clinical information associated with these samples was not publicly available.

All tissue samples were snap-frozen and embedded in optimal cutting temperature (OCT) compound. The embedded tissues were sectioned into 10-μm slices and mounted onto 10x Genomics Visium slides for spatial transcriptomic profiling. RNA libraries were subsequently prepared and sequenced on an Illumina NovaSeq 6000 platform for downstream analysis.

### Sequencing, Data Processing, and Filtering

We further processed the data using Space Ranger, a software tool developed by 10x Genomics for spatial transcriptomic analysis. Space Ranger uses raw RNA sequencing files, an image of the tissue section on the slide, slide barcode information, and a reference genome to generate files containing transcriptomic and spatial data.

We processed the ependymoma samples using Space Ranger v2.1.1 on Brigham Young University’s high-performance research computing cluster. We obtained all raw sequencing files from the GEO database associated with the study and downloaded the reference genome from the 10x Genomics website. In contrast, we did not process the diffuse midline glioma (DMG) samples ourselves using Space Ranger. Instead, we obtained the Space Ranger output files generated by the original research team using Space Ranger v1.1.0. Although the two software versions differ, they produce comparable outputs containing the information required for downstream analyses.

After obtaining the Space Ranger output files for each spatial transcriptomics dataset, we followed the data processing and filtering procedures described by Sibai et al. To reduce noise, we excluded genes with fewer than five unique molecular identifiers (UMIs). We then removed mitochondrial, ribosomal, and non-protein-coding genes to focus our analysis on nuclear protein-coding genes. Finally, we normalized gene expression values and generated Seurat objects containing normalized expression data adjusted for technical variation.

### Spatial Enhancement and Gene Expression Imputation with BayesSpace

Because each VISIUM spot has a diameter of ∼55 μm, each spot has the potential to contain multiple cells and thus mix signals from different cell types. To untangle these signals, we used BayesSpace (Zhao et al. 2021) to enhance cellular features in our samples. BayesSpace uses Bayesian statistics and information from nearby spots to refine the resolution of gene expression maps to the subspot level. By using BayesSpace to split each spot into six “subspots” with diameters of ∼20 μm, we better approximated the signals originating from individual cells in our samples.

### Assigning the Neoplastic and TME Compartments

We then categorized each subspot as cancerous or not. To facilitate annotation, we formatted our subspot-level enhanced gene expression data according to the Gene Cluster Text (GCT) file format specification. We then ran each sample’s subspot gene expression data into the ESTIMATE algorithm (Yoshihara et al. 2013), which infers stromal and immune cell infiltration based on gene expression profiles. By applying ESTIMATE, we derived immune and stromal scores for each subspot, reflecting the relative abundance of non-tumor cells. We then clustered subspots based on their immune and stromal scores using k-means clustering (k=5) to create a spectrum from cancer-dominated to TME-dominated regions. Clusters were re-labeled to represent this gradient, enabling clear differentiation between tumor and microenvironmental compartments. We are confident in these annotations, as Sibai et al. validated their accuracy with an expert human pathologist.

### Assigning Genes to Hallmarks

We used the same Cancer Hallmark transcriptional signatures developed by Sibai et al. (2025) to quantify hallmark activity. Sibai et al. developed transcriptional signatures for 13 of the 14 Cancer Hallmarks, explicitly excluding “Polymorphic Microbiomes” because their approach relied on human gene expression data and was not designed to capture microbial genetic material. To generate these signatures, the authors selected pathways from the Pathway Commons database and assembled sets of genes associated with each Hallmark. For example, the “Unlocking Phenotypic Plasticity” Hallmark was represented by gene sets from pathways including “Transcriptional regulation of pluripotent stem cells,” “Wnt signaling pathway,” and “Hedgehog signaling pathway.” Sibai et al. subsequently assessed the robustness of each Hallmark-associated gene list. Accordingly, our analysis used the 13 available Cancer Hallmark transcriptional signatures, consistent with the framework established by Sibai et al. For a complete description of the signature development and validation methods, readers are referred to Sibai et al. (2025).

### Scoring Cancer Hallmarks and Differential Gene Expression Analysis

After compiling gene transcription profiles associated with each Cancer Hallmark, we quantified hallmark activity in each subspot using the AddModuleScore_UCell() function from the UCell package. UCell calculates a gene signature enrichment score for each subspot by ranking genes according to their expression levels and evaluating the relative ranks of genes within each hallmark signature. These ranks are used to calculate a normalized enrichment score based on a Mann–Whitney U statistic, yielding scores between 0 and 1, with higher scores indicating greater enrichment of the corresponding hallmark signature. Because UCell evaluates gene expression rankings independently within each subspot,its scores are less sensitive to differences in dataset composition and are more robust to variation across samples and batches.

This approach differs from that of Sibai et al., who used Seurat’s AddModuleScore() function to quantify gene signature activity. Unlike UCell, AddModuleScore() calculates the average expression of genes within a signature and subtracts the average expression of a set of control genes selected to have comparable baseline expression levels. Although both methods quantify the relative activity of biological programs, they differ in how enrichment is measured: UCell relies on within-subspot gene expression rankings, whereas AddModuleScore() relies on average expression relative to matched control genes. We therefore used UCell to assess Cancer Hallmark activity across subspots, providing a rank-based scoring approach that is less dependent on the overall expression distribution and cellular composition of the dataset.

With Hallmark scores computed at the subspot level, we split each sample into the neoplastic and TME compartments, calculated median hallmark scores within each compartment, and aggregated these median values across all samples to determine which compartment each Cancer Hallmark was most highly expressed in. We compiled these compartment median values into a heatmap and scaled by row (by Cancer Hallmark) to emphasize compartmental differences within each hallmark. Our first such heatmap revealed one sample to contain extreme outlier hallmark scores. To clarify and emphasize the overall distribution of Cancer Hallmarks between compartments, we removed this sample from our analysis. The results from our analysis remained mostly unchanged after removing the sample (**Fig. S1A**).

To examine which genes in each Hallmark were contributing to Hallmarks’ compartmentalization, we performed differential gene expression analysis. For each sample, and each gene contained in a certain hallmark, we computed the mean gene expression values in the neoplastic compartment and TME compartment. We then aggregated these means across all samples and performed paired Wilcoxon signed-rank tests. For each gene, we also computed the log₂ fold change (neoplastic mean gene expression divided by TME mean gene expression). Genes with an absolute log₂ fold change greater than 1 and a p-value less than 0.05 were considered significantly differentially expressed. These values were visualized in volcano plots, with log₂ fold change on the x-axis and the negative log₁₀-transformed p-value on the y-axis. Genes meeting significance criteria were highlighted and labeled. To compare our findings with adult brain tumors, we obtained similar plots from Sibai et al., restricted to their adult brain tumor samples (**Fig. S3**).

### Spatial Autocorrelation of Cancer Hallmark Activities

To assess the spatial autocorrelation of hallmark activity scores across tissue sections, Sibai et al. adapted the CorSpatialGenes() function from the STutility package to accommodate enhanced-resolution Visium spatial transcriptomics data. Specifically, they implemented a modified version of the function that incorporates corrected subspot coordinates to more accurately define local spatial neighborhoods. For each subspot, a spatial network was constructed by identifying its 10 nearest neighbors: 5 within the same spot (intra-spot) and 5 from adjacent spots (inter-spot). Edge weights were assigned inversely proportional to the corrected Euclidean distances between neighboring subspots. Spatial autocorrelation was then calculated as the Pearson correlation between each hallmark’s subspot expression values and its corresponding spatial lag vector. A spatial lag vector is a weighted average of the expression values from neighboring locations surrounding a given observation, calculated using spatial weights. In this analysis, it represents the local expression pattern surrounding each subspot and is used in the code to quantify the similarity between a subspot’s expression and that of its spatial neighbors. Using this modified function, we calculated the spatial autocorrelation of each cancer hallmark across entire tissue samples, as well as separately within the neoplastic and tumor microenvironment (TME) compartments.

### Volcano Plots

To further characterize differences in cancer hallmark activity between the neoplastic and tumor microenvironment (TME) compartments, we performed differential expression analyses for genes associated with each of the 13 cancer hallmarks. For each gene, mean expression values were calculated separately across the neoplastic and TME compartments. Differential expression was assessed using a paired Wilcoxon signed-rank test, and the magnitude and direction of expression differences were quantified as the log₂ fold change (neoplastic/TME). Genes were considered significantly differentially expressed if they met both criteria: a p-value below 0.05 and an absolute log₂ fold change greater than 1. Results were visualized using volcano plots, with significantly differentially expressed genes highlighted and labeled. Finally, the number of significantly differentially expressed genes was quantified for each hallmark and summarized in a bar plot to compare the extent of differential expression across hallmarks (**Fig. S5**).

### Modeling Cross-Compartment Inter-Hallmark Interactions with Random Forests

Prior to modeling cross-compartment hallmark interactions, we ensured all hallmark scores were strictly positive by subtracting the minimum value of each hallmark across the tissue and adding a small constant (0.001). This normalization step preserves relative differences while avoiding negative values that could distort downstream distance-weighted calculations.

For each subspot in the target compartment, we calculated a radar score for each predictor hallmark as a distance-weighted sum of that hallmark’s expression in surrounding predictor subspots. The inverse distance was used as a weight to prioritize closer predictor subspots, reflecting the assumption that spatial influence diminishes with distance. In parallel, we extracted the local target-specific hallmark scores directly from each target subspot. This yielded a matrix in which each target subspot is annotated both with intrinsic hallmark scores and extrinsic predictor radar scores.

For each sample, we aggregated the predictor radar scores and target hallmark scores to train Random Forest regression models. These models were constructed to predict the spatial activity of individual target hallmarks based on the activity levels of multiple predictor hallmarks. For each target hallmark, we assembled a dataset consisting of its corresponding hallmark scores across subspots as the dependent variable and the associated predictor hallmark radar scores as independent variables. This process was repeated for each sample individually, allowing us to examine sample-specific relationships between neoplastic and microenvironmental programs.

To assess the predictive relationships between neoplastic and TME hallmark scores, we performed two machine learning experiments:

1. Predicting neoplastic hallmarks from TME hallmark radar scores: For each of the three individual neoplastic hallmarks, we trained a model using the six TME hallmark radar scores as input features. This was done independently for each of the 17 samples, resulting in a total of 3 targets × 17 samples = 51 models.
2. Predicting TME hallmarks from neoplastic hallmark radar scores: For each of the six individual TME hallmarks, we trained a model using the three neoplastic hallmark radar scores as input features. This yielded a total of 6 targets × 17 samples = 102 models.

For each target hallmark, we split our data into a training set (80%) and a testing set (20%) using stratified sampling to preserve the distribution of the response variable. We then trained a Random Forest model using the ranger package with 500 trees and impurity-based feature importance, allowing the model to identify which predictor hallmarks most influence the spatial distribution of the target hallmarks. We then used the treeshap package to unify the trained model and apply TreeSHAP, which calculates SHAP values – interpretable, sample-level, estimates of how each predictor contributes to the model’s output. A SHAP value measures how much a single feature contributed to a model’s prediction for a specific input, by comparing the prediction to the average model output across the entire dataset (the baseline), and calculating how the feature shifts the prediction when added to all possible combinations of the other features.

We used these SHAP values to further calculate relative importance and spatial dependency. We calculated relative importance for a single model by first taking the absolute value of the SHAP scores for all predictors, computing the mean SHAP value for each predictor, applying min-max scaling to these means, and then normalizing them by dividing each scaled value by the sum of all scaled values. The resulting normalized values represent the relative importance of each predictor within that model, reflecting each predictor’s proportional contribution to the overall prediction across all features. Whereas relative importance quantifies the strength of hallmark interactions across compartments, spatial dependence quantifies how much these interactions depend on spatial proximity. To calculate this spatial dependency, we computed the correlation between target subspots’ radar scores and their corresponding SHAP values indicating their effect on the model’s prediction. Because radar scores account for spatial proximity, a high correlation between radar scores and SHAP values suggests that the hallmark interaction is highly dependent on spatial proximity. This approach enables quantification and interpretation of the spatial dependencies between neoplastic and TME programs across samples, highlighting potential spatial cross-talk between the tumor and microenvironment.

### Calculating Tumor Purity

We calculated tumor purity by assessing the spatial organization of each subspot’s ESTIMATE score relative to its neighboring subspots. Specifically, we calculated the overall spatial autocorrelation of subspot ESTIMATE scores within each sample; a score we refer to as a Spatial Coherence Score (SCS). Higher spatial autocorrelation indicates greater spatial clustering of ESTIMATE scores, whereas lower autocorrelation indicates a more spatially random distribution of scores throughout the sample. We used this spatial autocorrelation measure as a proxy for the spatial organization of neoplastic and tumor microenvironment (TME) compartments.

After calculating this measure for each sample, we examined its correlation with the average expression of each neoplastic Cancer Hallmark gene. For each sample, we calculated the average expression of each selected gene across subspots classified as neoplastic (ESTIMATE score 1 or 2). This analysis was performed separately for genes associated with the three strongest pediatric neoplastic hallmarks. We then calculated Pearson correlation coefficients across samples between each gene’s average expression and the corresponding sample’s spatial autocorrelation-based tumor-purity measure. Genes with missing correlation values were excluded from subsequent analyses.

To further characterize the relationship between tumor purity and cancer-associated genes, we used the OncoKB Cancer Gene List (downloaded June 10, 2025) to annotate genes according to their oncogene and tumor suppressor status. We examined correlations for all genes and separately for genes included in the OncoKB list. For visualization, we identified genes with strong positive or negative correlations, using absolute correlation thresholds of greater than 0.5 for the full gene set and greater than 0.3 for the OncoKB-annotated subset. Finally, among genes included in the OncoKB list, we used chi-square tests to assess whether the direction of correlation (positive versus negative) was associated with oncogene status.

### Cell Type

To characterize the cellular composition of the pediatric TME relative to the neoplastic compartment, we sought to identify the cell types associated with each spatial compartment. We leveraged PanglaoDB (Franzén et al., 2019), a publicly available database that curates cell type-specific gene expression signatures derived from single-cell RNA sequencing (scRNA-seq) studies. Using marker gene signatures representing 42 cell types across three broad cellular groups (nervous system, immune system, and vasculature), we calculated “cell type enrichment scores” for each spatial subspot. This approach allowed us to characterize the likely cellular identity of individual subspots based on their transcriptional similarity to established cell type-specific expression profiles.

To characterize the cellular composition of the pediatric TME relative to the neoplastic compartment, we sought to identify the cell types associated with each spatial compartment. We leveraged PanglaoDB (Franzén et al., 2019), a publicly available database that curates cell type-specific gene expression signatures derived from single-cell RNA sequencing (scRNA-seq) studies. Using marker gene signatures representing 42 cell types across three broad cellular groups (nervous system, immune system, and vasculature), we calculated cell type enrichment scores for each spatial subspot. This approach allowed us to characterize the likely cellular identity of individual subspots based on their transcriptional similarity to established cell type-specific expression profiles.

Human-specific canonical marker genes were extracted from the PanglaoDB marker gene database (March 27, 2020 release). Marker genes were grouped according to their annotated cell types and broad cellular categories, including the nervous system (brain), immune system, and vasculature. To ensure sufficient marker representation, cell type-specific gene sets containing at least 10 canonical marker genes were retained for downstream analysis.

Cell type enrichment scores were calculated using the AddModuleScore function in the Seurat R package. An enhanced subspot-level gene expression matrix generated following gene expression imputation was used as input. The expression data were normalized and scaled using Seurat’s NormalizeData and ScaleData functions, respectively. For each cell type-specific marker gene set, enrichment scores were calculated as the average expression of the marker genes relative to expression-matched control genes selected from the background gene pool. The scoring procedure used 24 expression bins and 100 control genes per marker gene, with a random seed of 200 to ensure reproducibility. Higher enrichment scores indicated greater relative expression of the corresponding cell type-specific marker signature within a given subspot.

To characterize cell type enrichment across spatial compartments, subspot-level enrichment scores were aggregated according to their corresponding ESTIMATE-derived compartment scores. Mean enrichment scores were calculated for each compartment within each sample. These sample-level averages were subsequently integrated across the cohort to enable comparisons of cell type-associated transcriptional signatures across samples and tumor types. Samples were annotated according to their originating study and cancer type. The resulting cell type enrichment profiles were visualized using hierarchical clustering heatmaps. Prior to visualization, enrichment scores were standardized across samples for each cell type (row-wise scaling). Hierarchical clustering was performed using correlation-based distances and average linkage for both cell types and samples.

To investigate associations between cellular signatures and biological pathway activity, cell type enrichment scores were compared with previously calculated hallmark pathway activity scores at the subspot level. Pearson correlation coefficients were calculated between each hallmark pathway score and each cell type enrichment score across subspots within individual samples. The resulting correlation matrices were subsequently integrated across samples by calculating the arithmetic mean correlation coefficient for each cell type–hallmark pathway pair. These mean correlation coefficients were visualized using hierarchical clustering heatmaps, with correlation values standardized across cell types for each hallmark pathway (column-wise scaling). Hierarchical clustering was performed using correlation-based distances and average linkage to identify patterns of association between cell type signatures and hallmark pathway activity across the cohort.

## Supplementary Figure Captions

**Supplemental Figure 1: Metrics of the gene used in the cancer hallmark scoring.**

**(A)** A barplot indicates the frequency (y-axis) a gene is part of multiple hallmarks, e.g., ∼500 genes are observed in 2 different hallmark genesets. (

**B)** A barplot displays the number of genes (y-axis) per hallmark (x-axis)

**C)** A barplot displays the number of pathways that comprise each hallmark. Of note, when Sabai et al. restricted their analysis to a random set of half of genes per hallmark, the same results and separation was observed in their analyses. This subsetting of genes was not applied in our study.

**Supplemental Figure 2: Heatmap of the Cancer Hallmarks by ESTIMATE-assigned compartments.**

**A)** Similar to **Figure 2A** in the manuscript, this figure highlights clear separation of Hallmark activity between tumor and normal compartments, but also highlights one outlier expression sample that was removed from our main analysis sample (SRR17814899). Removal of this sample did not meaningfully alter the results of our analysis. An unsupervised clustered heatmap of spatial transcriptomics spots by Cancer Hallmark Score (dendrogram removed). Columns indicate sample-compartments as defined by tumor purity (purple is more neoplastic and yellow is more TME). Two additional column annotations include study and pediatric brain cancer type. Rows indicate Cancer Hallmark scores as estimated by gene sets and labels with Hallmark symbols and names.

**B)** Similar to **Figure 2A** but restricted to Glioblastoma samples from Sabai et al. manuscript.

**Supplemental Figure 3: Differential Expression Analysis Between Adult GBM TME and Neoplastic Compartments (Compare to Figure 3)**

Corresponding Volcano Plots comparing mean adult GBM Cancer Hallmark scores between neoplastic and TME compartments across samples. Compare with Figure 3. Log₂ fold change (neoplastic scores divided by TME scores on the x-axis) is plotted against statistical significance (-log_10_ p-value on the y-axis). Thus, values to the left of zero are more expressed in the TME, and points to the right of zero are more expressed in the neoplastic compartments. Red points indicate significant genes. **A)** Sustaining Proliferative Signaling**; B)** Evading Growth Suppressors; **C)** Avoiding Immune Destruction**; D)** Tumor-Promoting Inflammation**; E)** Activating Invasion and Metastasis**; F)** Inducing Angiogenesis**; G)** Resisting Cell Death; **H)** Deregulating Cellular Energetics; **I)** Unlocking Phenotypic Plasticity; **J)** Enabling Replicative Immortality; **K)** Genome Instability and Mutation**; L)** Senescent Cells; and **M)** Nonmutational Epigenetic Reprogramming.

**Supplemental Figure 4: Spatial Dependencies and Model-Predictor Importances Across Tumor Compartments restricted to fewer hallmarks in the model. (Compare to Figure 4)**

**A)** Analytical workflow to establish compartment-specific hallmark scoring for cross compartment modeling. <u>Step 0:</u> An illustration shows sample transcriptomic subspots divided into two compartments, neoplastic and TME. <u>Step 1:</u> Hallmark expression scores are highlighted in one compartment. <u>Step 2:</u> A visual representation of projecting one compartment onto another. d1 and d2 illustrate that each compartment projection is weighted by the distance between subspots. These projections will be used as predictors in our model. <u>Step 3:</u> Shows that the targets of the predictors are the original hallmark scores. <u>Step 4:</u> Illustrates a repeat of step 2 for each hallmark to populate the features of the model. <u>Step 5:</u> Illustrates how each hallmark from the opposing compartment was included in the model to predict one Hallmark. <u>Step 6:</u> Evaluate pairs of the hallmarks from the model for possible cross compartment Hallmark interactions.

**B)** A bar chart shows the summed relative importance of TME radar predictors, i.e., features of the model.

**C)** A bar chart shows the summed relative importance of neoplastic radar predictors.

**D)** A heatmap shows TME predictor-importance across neoplastic hallmarks. Predictor-Hallmarks are on the x-axis, and the predicted-Hallmarks are on the y-axis.

**E)** A heatmap indicates Neoplastic predictor-importance heatmap across TME hallmarks. Again, Predictor-Hallmarks are on the x-axis, and the predicted-Hallmarks are on the y-axis.

**F)** A ridgeplot displays spatial dependency distributions for TME hallmarks.

**G)** A ridgeplot displays spatial dependency distributions for neoplastic hallmarks.

**H)** A heatmap shows Pearson correlation across subspots between the model’s radar score and its SHAP value in the target-hallmark model. Red values indicate Hallmarks that tend to be active near one another. Blue values tend to be distant from one another. TME predictors are on the x-axis, and the predicted neoplastic relationships are on the y-axis.

**I)** Same as panel H but with swapped TME and neoplastic axes.

**Supplemental Figure 5: Counts of Significantly Differentially Expressed Genes in Pediatric Samples**

Expanding on Figure 3. The number of genes that are significantly differentially expressed between the neoplastic and TME compartments in pediatric samples per Cancer Hallmark.

**Supplemental figure 6-22: Hallmark composition for each sample.** For each sample, a comprehensive suite of images is provided. First, an H&E stain. Second, ESTIMATE cluster assignments (purple = neoplastic, yellow = TME). Third, a max hallmark spot assignment score to see the max (normalized) hallmark-score per spot. And finally as a series of Hallmark scores for each tumor.

## Supplemental Table

**Supplemental Table 1:** Metadata for the samples used.

| Tissue site | Data Origin | Original ID | Embedding method | Number of spots | Total number of detected genes | Median genes per spot | Minimum number of features threshold used to filter spots | WHO Grade | Age (yrs) |
|---|---|---|---|---|---|---|---|---|---|
| Ependymoma | PMID: 36215273 | SRR17814893_SR | OCT | 1264 | 22824 | 4102.5 | 250 | III | 5 |
| Ependymoma | PMID: 36215273 | SRR17814894_SR | OCT | 3618 | 23755 | 3425.5 | 250 | III | 0.9 |
| Ependymoma | PMID: 36215273 | SRR17814895_SR | OCT | 2963 | 23575 | 3192 | 250 | III | 1.9 |
| Ependymoma | PMID: 36215273 | SRR17814896_SR | OCT | 2049 | 22748 | 3301 | 250 | III | 3 |
| Ependymoma | PMID: 36215273 | SRR17814897_SR | OCT | 2750 | 23374 | 2843.5 | 250 | II | 6 |
| Ependymoma | PMID: 36215273 | SRR17814898_SR | OCT | 3156 | 22451 | 2751 | 250 | III | 2 |
| Ependymoma | PMID: 36215273 | SRR17814899_SR | OCT | 1841 | 23252 | 4700 | 250 | III | 1.9 |
| Ependymoma | PMID: 36215273 | SRR17814900_SR | OCT | 3026 | 23815 | 3497.5 | 250 | III | 0.6 |
| Ependymoma | PMID: 36215273 | SRR17814901_SR | OCT | 1610 | 23837 | 4803 | 250 | III | 6 |
| Ependymoma | PMID: 36215273 | SRR17814902_SR | OCT | 2704 | 23844 | 3453.5 | 250 | III | 5 |
| Ependymoma | PMID: 36215273 | SRR17814903_SR | OCT | 3059 | 24211 | 3068 | 250 | III | 0.9 |
| Ependymoma | PMID: 36215273 | SRR17814904_SR | OCT | 978 | 22952 | 6391.5 | 250 | II | 2 |
| Ependymoma | PMID: 36215273 | SRR17814905_SR | OCT | 1959 | 22870 | 3815 | 250 | II | 2 |
| Ependymoma | PMID: 36215273 | SRR17814906_SR | OCT | 2237 | 23596 | 4500 | 250 | III | 0.5 |
| Diffuse Midline Glioma | GEO Dataset: GSE194329 | DMG2 | OCT | 2305 | 23797 | 5161 | 250 | NA | <18 |
| Diffuse Midline Glioma | GEO Dataset: GSE194329 | DMG3 | OCT | 1063 | 23375 | 7221 | 250 | NA | <18 |
| Diffuse Midline Glioma | GEO Dataset: GSE194329 | DMG4 | OCT | 1056 | 23003 | 5220.5 | 250 | NA | <18 |
| Diffuse Midline Glioma | GEO Dataset: GSE194329 | DMG5 | OCT | 2128 | 23520 | 4075 | 250 | NA | <18 |

## REFERENCES

Aran, Dvir, Marina Sirota, and Atul J. Butte. 2015. “Systematic Pan-Cancer Analysis of Tumour Purity.” Nature Communications 6 (1): 8971.

Bockmayr, Michael, Malte Mohme, Frederick Klauschen, et al. 2018. “Subgroup-Specific Immune and Stromal Microenvironment in Medulloblastoma.” Oncoimmunology 7 (9): e1462430.

Chakravarty, Debyani, Jianjiong Gao, Sarah M. Phillips, et al. 2017. “OncoKB: A Precision Oncology Knowledge Base.” JCO Precision Oncology 2017 (July).

Del Bigio, Marc R. 2010. “Ependymal Cells: Biology and Pathology.” Acta Neuropathologica 119 (1): 55–73.

Formentin, Cleiton, Andrei Fernandes Joaquim, and Enrico Ghizoni. 2023. “Posterior Fossa Tumors in Children: Current Insights.” European Journal of Pediatrics 182 (11): 4833–4850.

Fu, Rui, Gregory A. Norris, Nicholas Willard, et al. 2023. “Spatial Transcriptomic Analysis Delineates Epithelial and Mesenchymal Subpopulations and Transition Stages in Childhood Ependymoma.” Neuro-Oncology 25 (4): 786–798.

Gates, Evan D. H., Jie Yang, Kazutaka Fukumura, et al. 2019. “Spatial Distance Correlates with Genetic Distance in Diffuse Glioma.” Frontiers in Oncology 9 (July): 676.

Getis, Arthur. 2007. “Reflections on Spatial Autocorrelation.” Regional Science and Urban Economics 37 (4): 491–496.

Gonzalez Castro, L. Nicolas, Ilon Liu, and Mariella Filbin. 2023. “Characterizing the Biology of Primary Brain Tumors and Their Microenvironment via Single-Cell Profiling Methods.” Neuro-Oncology 25 (2): 234–247.

Greenwald, Alissa C., Noam Galili Darnell, Rouven Hoefflin, et al. 2024. “Integrative Spatial Analysis Reveals a Multi-Layered Organization of Glioblastoma.” Cell 187 (10): 2485–2501.e26.

Griesinger, Andrea M., Diane K. Birks, Andrew M. Donson, et al. 2013. “Characterization of Distinct Immunophenotypes across Pediatric Brain Tumor Types.” Journal of Immunology (Baltimore, Md.: 1950) 191 (9): 4880–4888.

Gröbner, Susanne N., Barbara C. Worst, Joachim Weischenfeldt, et al. 2018. “The Landscape of Genomic Alterations across Childhood Cancers.” Nature 555 (7696): 321–327.

Gurney, J. 1999. “CNS and Miscellaneous Intracranial and Intraspinal Neoplasms.” Cancer Incidence and Survival among Children and Adolescents : United States SEER Program 1975-1995, 51–63.

Hanahan, Douglas. 2022. “Hallmarks of Cancer: New Dimensions.” Cancer Discovery 12 (1): 31–46.

Hanahan, Douglas, and Robert A. Weinberg. 2011. “Hallmarks of Cancer: The next Generation.” Cell 144 (5): 646–674.

Hanahan, D., and R. A. Weinberg. 2000. “The Hallmarks of Cancer.” Cell 100 (1): 57–70.

Jessa, Selin, Alexis Blanchet-Cohen, Brian Krug, et al. 2019. “Stalled Developmental Programs at the Root of Pediatric Brain Tumors.” Nature Genetics 51 (12): 1702–1713.

Kline, Nancy E., and Nicole Sevier. 2003. “Solid Tumors in Children.” Journal of Pediatric Nursing 18 (2): 96–102.

Lawlor, Elizabeth R., and Carol J. Thiele. 2012. “Epigenetic Changes in Pediatric Solid Tumors: Promising New Targets.” Clinical Cancer Research: An Official Journal of the American Association for Cancer Research 18 (10): 2768–2779.

Lawrence, Michael S., Petar Stojanov, Paz Polak, et al. 2013. “Mutational Heterogeneity in Cancer and the Search for New Cancer-Associated Genes.” Nature 499 (7457): 214–218.

Lee-Six, Henry, Taryn D. Treger, Manas Dave, et al. 2025. “High Resolution Clonal Architecture of Hypomutated Wilms Tumours.” Nature Communications 16 (1): 4647.

Lewis, Sabrina M., Marie-Liesse Asselin-Labat, Quan Nguyen, et al. 2021. “Spatial Omics and Multiplexed Imaging to Explore Cancer Biology.” Nature Methods 18 (9): 997–1012.

Lieberman, Nicole A. P., Kole DeGolier, Heather M. Kovar, et al. 2019. “Characterization of the Immune Microenvironment of Diffuse Intrinsic Pontine Glioma: Implications for Development of Immunotherapy.” Neuro-Oncology 21 (1): 83–94.

Lin, Grant L., Surya Nagaraja, Mariella G. Filbin, Mario L. Suvà, Hannes Vogel, and Michelle Monje. 2018. “Non-Inflammatory Tumor Microenvironment of Diffuse Intrinsic Pontine Glioma.” Acta Neuropathologica Communications 6 (1): 51.

Majzner, Robbie G., Jason S. Simon, Joseph F. Grosso, et al. 2017. “Assessment of Programmed Death-Ligand 1 Expression and Tumor-Associated Immune Cells in Pediatric Cancer Tissues.” Cancer 123 (19): 3807–3815.

Ma, Xiaotu, Yu Liu, Yanling Liu, et al. 2018. “Pan-Cancer Genome and Transcriptome Analyses of 1,699 Paediatric Leukaemias and Solid Tumours.” Nature 555 (7696): 371–376.

Nikbakht, Hamid, Eshini Panditharatna, Leonie G. Mikael, et al. 2016. “Spatial and Temporal Homogeneity of Driver Mutations in Diffuse Intrinsic Pontine Glioma.” Nature Communications 7 (1): 11185.

Park, Jin A., Jeyeon Im, and Sung-Min Hwang. 2026. “Decoding T Cell Exhaustion in the Tumour Microenvironment.” Experimental & Molecular Medicine 58 (8): 2590–2602.

Patel, Jenny P., Susan E. Spiller, and Elizabeth D. Barker. 2021. “Drug Penetration in Pediatric Brain Tumors: Challenges and Opportunities.” Pediatric Blood & Cancer 68 (6): e28983.

Rahal, Zahraa, Farah Abdulhai, Humam Kadara, and Raya Saab. 2018. “Genomics of Adult and Pediatric Solid Tumors.” American Journal of Cancer Research 8 (8): 1356–1386.

Ren, Yanming, Zongyao Huang, Lingling Zhou, et al. 2023. “Spatial Transcriptomics Reveals Niche-Specific Enrichment and Vulnerabilities of Radial Glial Stem-like Cells in Malignant Gliomas.” Nature Communications 14 (1): 1028.

Rinaldi, Gianmarco, Matteo Rossi, and Sarah-Maria Fendt. 2018. “Metabolic Interactions in Cancer: Cellular Metabolism at the Interface between the Microenvironment, the Cancer Cell Phenotype and the Epigenetic Landscape.” Wiley Interdisciplinary Reviews. Systems Biology and Medicine 10 (1): e1397.

Sibai, Mustafa, Sergi Cervilla, Daniela Grases, et al. 2025. “The Spatial Landscape of Cancer Hallmarks Reveals Patterns of Tumor Ecological Dynamics and Drug Sensitivity.” Cell Reports 44 (2): 115229.

Ståhl, Patrik L., Fredrik Salmén, Sanja Vickovic, et al. 2016. “Visualization and Analysis of Gene Expression in Tissue Sections by Spatial Transcriptomics.” Science 353 (6294): 78–82.

Strobl, Carolin, Anne-Laure Boulesteix, Achim Zeileis, and Torsten Hothorn. 2007. “Bias in Random Forest Variable Importance Measures: Illustrations, Sources and a Solution.” BMC Bioinformatics 8 (1): 25.

Tickle, Timothy, Itay Tirosh, Christophe Georgescu, Maxwell Brown, Brian Haas, and Others. 2019. “InferCNV of the Trinity CTAT Project.” Klarman Cell Observatory, Broad Institute of MIT and Harvard.

Torp, Sverre Helge, Ole Solheim, and Anne Jarstein Skjulsvik. 2022. “The WHO 2021 Classification of Central Nervous System Tumours: A Practical Update on What Neurosurgeons Need to Know-a Minireview.” Acta Neurochirurgica 164 (9): 2453–2464.

Vignali, Paolo D. A., Kristin DePeaux, Mclane J. Watson, et al. 2023. “Hypoxia Drives CD39-Dependent Suppressor Function in Exhausted T Cells to Limit Antitumor Immunity.” Nature Immunology 24 (2): 267–279.

Yang, Jian, Jiajia Wang, Shuaiwei Tian, et al. 2021. “An Integrated Analysis of Tumor Purity of Common Central Nervous System Tumors in Children Based on Machine Learning Methods.” Frontiers in Genetics 12 (December): 707802.

Yan, Junli, Jianming Jiang, Ching Aeng Lim, Qiang Wu, Huck-Hui Ng, and Keh-Chuang Chin. 2007. “BLIMP1 Regulates Cell Growth through Repression of p53 Transcription.” Proceedings of the National Academy of Sciences of the United States of America 104 (6): 1841–1846.

Yoshihara, Kosuke, Maria Shahmoradgoli, Emmanuel Martínez, et al. 2013. “Inferring Tumour Purity and Stromal and Immune Cell Admixture from Expression Data.” Nature Communications 4 (1): 2612.

Zhao, Edward, Matthew R. Stone, Xing Ren, et al. 2021. “Spatial Transcriptomics at Subspot Resolution with BayesSpace.” Nature Biotechnology 39 (11): 1375–1384.

