## Supplemental figures for "Spatial Transcriptomics Reveals Age-Specific Cancer Hallmark Architecture in Pediatric and Adult Brain Tumors"

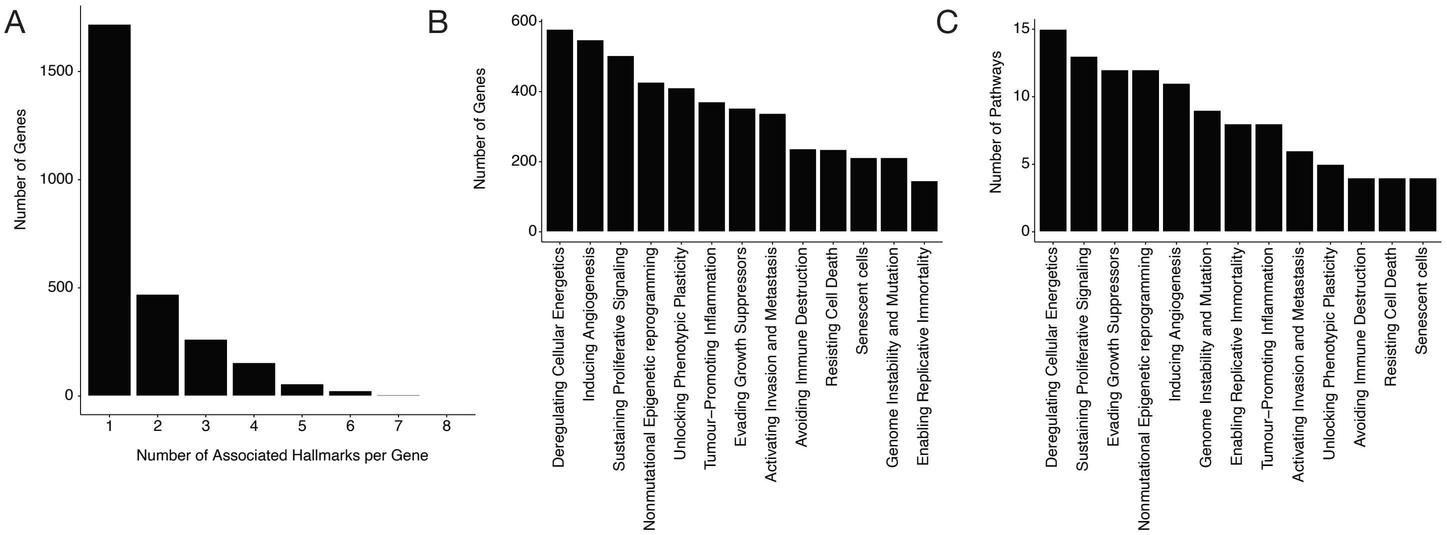

A
Number of Genes
Number of Associated Hallmarks per Gene
B
Number of Genes
Deregulating Cellular Energetics
Inducing Angiogenesis
Sustaining Proliferative Signaling
Nonmutational Epigenetic reprogramming
Unlocking Phenotypic Plasticity
Tumour−Promoting Inflammation
Evading Growth Suppressors
Activating Invasion and Metastasis
Avoiding Immune Destruction
Resisting Cell Death
Senescent cells
Genome Instability and Mutation
Enabling Replicative Immortality
C
Number of Pathways
Deregulating Cellular Energetics
Sustaining Proliferative Signaling
Evading Growth Suppressors
Nonmutational Epigenetic reprogramming
Inducing Angiogenesis
Genome Instability and Mutation
Enabling Replicative Immortality
Tumour−Promoting Inflammation
Activating Invasion and Metastasis
Unlocking Phenotypic Plasticity
Avoiding Immune Destruction
Resisting Cell Death
Senescent cells


Figure S1

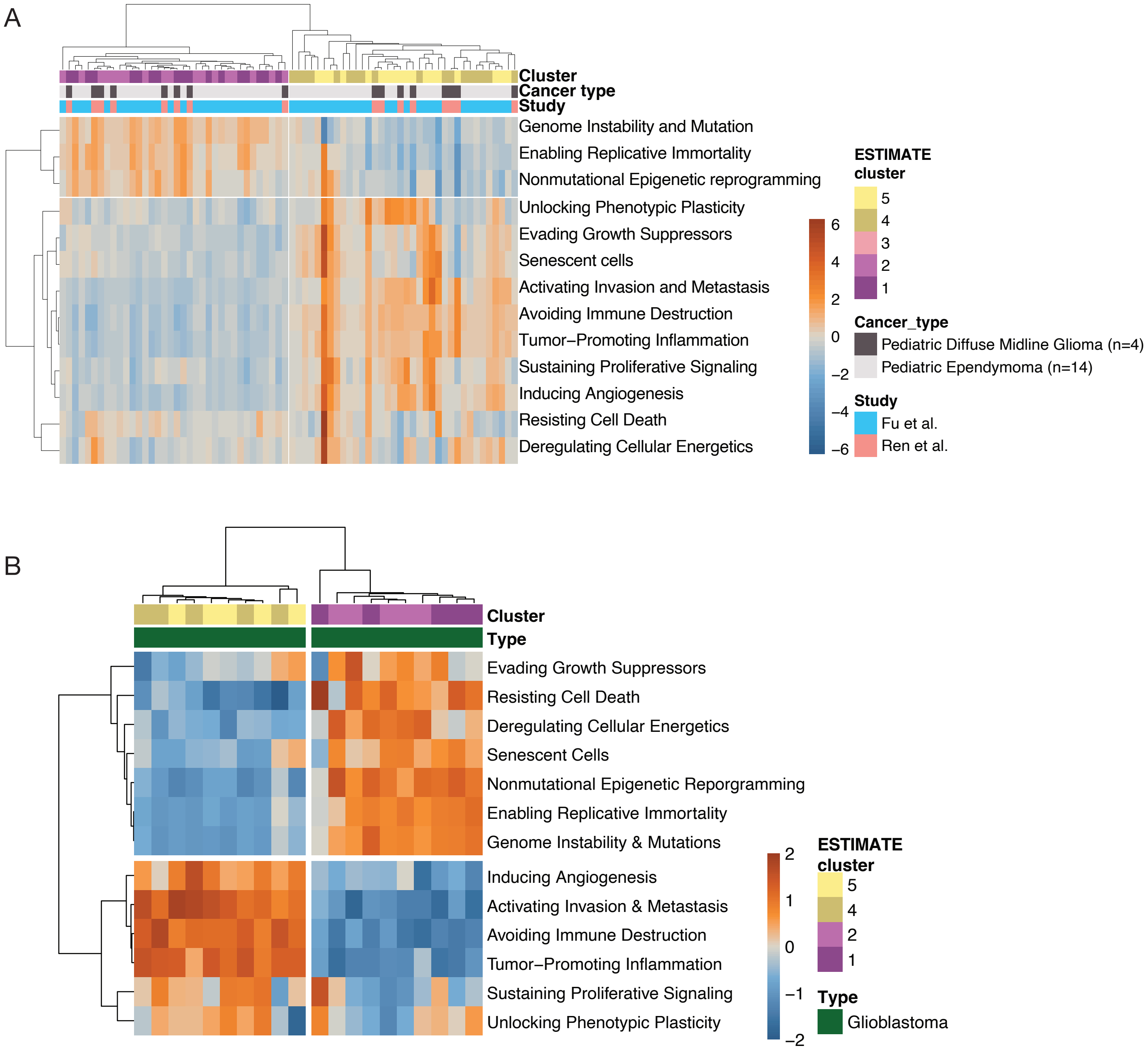
A
Cluster
Cancer type
Study
Genome Instability and Mutation
Enabling Replicative Immortality
Nonmutational Epigenetic reprogramming
Unlocking Phenotypic Plasticity
Evading Growth Suppressors
Senescent cells
Activating Invasion and Metastasis
Avoiding Immune Destruction
Tumor–Promoting Inflammation
Sustaining Proliferative Signaling
Inducing Angiogenesis
Resisting Cell Death
Deregulating Cellular Energetics
6
4
2
0
−2
−4
−6
ESTIMATE
cluster
5
4
3
2
1
Cancer_type
Pediatric Diffuse Midline Glioma (n=4)
Pediatric Ependymoma (n=14)
Study
Fu et al.
Ren et al.
B
Cluster
Type
Evading Growth Suppressors
Resisting Cell Death
Deregulating Cellular Energetics
Senescent Cells
Nonmutational Epigenetic Reporgramming
Enabling Replicative Immortality
Genome Instability & Mutations
Inducing Angiogenesis
Activating Invasion & Metastasis
Avoiding Immune Destruction
Tumor–Promoting Inflammation
Sustaining Proliferative Signaling
Unlocking Phenotypic Plasticity
2
1
0
−1
−2
ESTIMATE
cluster
5
4
2
1
Type
Glioblastoma

Figure S2

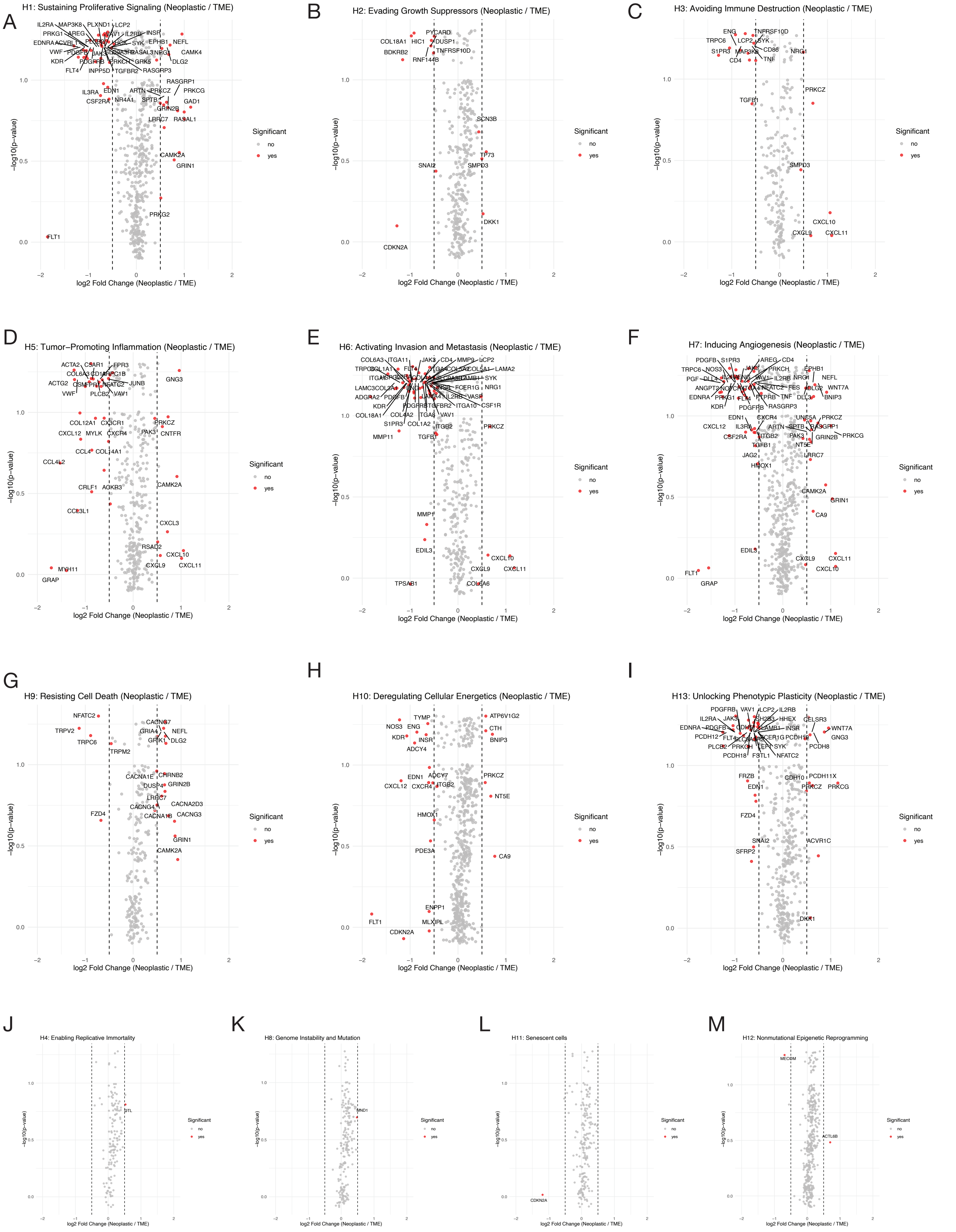
A
H1: Sustaining Proliferative Signaling (Neoplastic / TME)
B
H2: Evading Growth Suppressors (Neoplastic / TME)
C
H3: Avoiding Immune Destruction (Neoplastic / TME)
D
H5: Tumor−Promoting Inflammation (Neoplastic / TME)
E
H6: Activating Invasion and Metastasis (Neoplastic / TME)
F
H7: Inducing Angiogenesis (Neoplastic / TME)
G
H9: Resisting Cell Death (Neoplastic / TME)
H
H10: Deregulating Cellular Energetics (Neoplastic / TME)
I
H13: Unlocking Phenotypic Plasticity (Neoplastic / TME)
J
H4: Enabling Replicative Immortality
K
H8: Genome Instability and Mutation
L
H11: Senescent cells
M
H12: Nonmutational Epigenetic Reprogramming
−log10(p−value)
log2 Fold Change (Neoplastic / TME)
Significant
no
yes

Figure S3

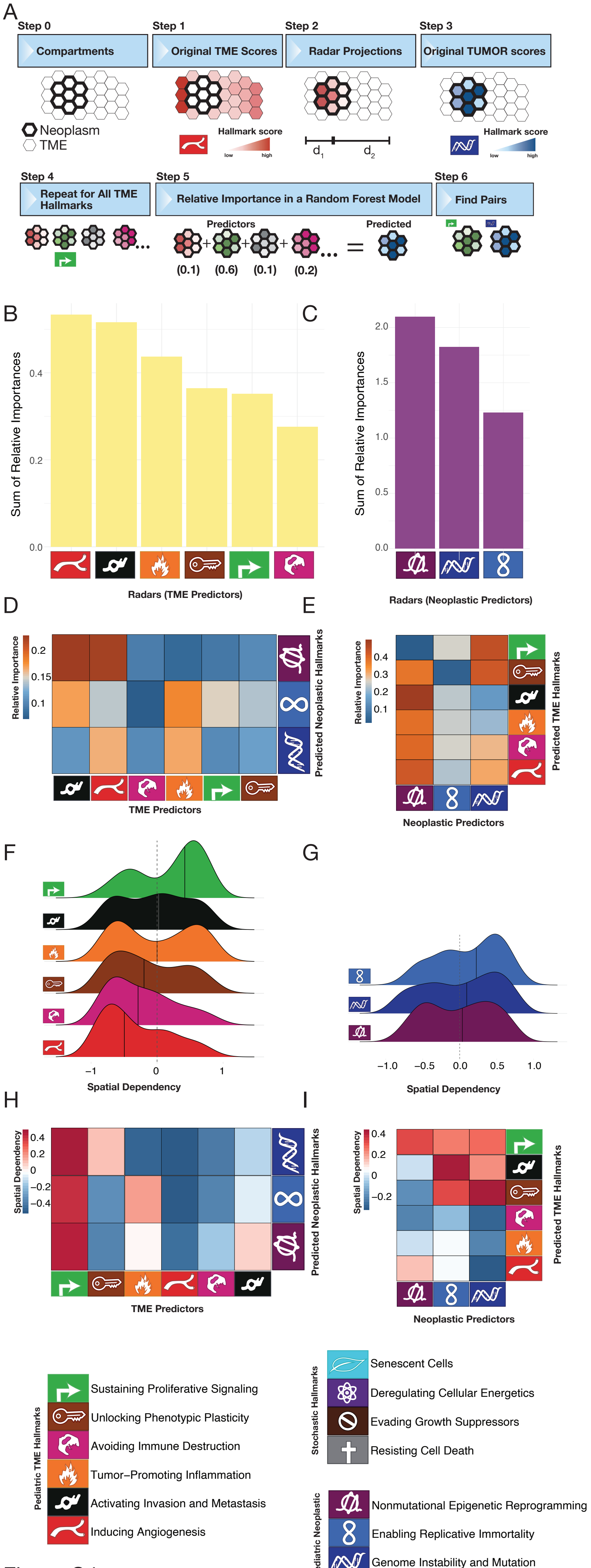
A
Step 0
Compartments
Neoplasm
TME
Step 1
Original TME Scores
Hallmark score
low
high
Step 2
Radar Projections
d1
d2
Step 3
Original TUMOR scores
Hallmark score
low
high
Step 4
Repeat for All TME Hallmarks
Step 5
Relative Importance in a Random Forest Model
Predictors
Predicted
(0.1) (0.6) (0.1) (0.2)
Step 6
Find Pairs
B
Sum of Relative Importances
0.0
0.2
0.4
Radars (TME Predictors)
C
Sum of Relative Importances
0.0
0.5
1.0
1.5
2.0
Radars (Neoplastic Predictors)
D
Relative Importance
0.2
0.15
0.1
TME Predictors
Predicted Neoplastic Hallmarks
E
Relative Importance
0.4
0.3
0.2
0.1
Neoplastic Predictors
Predicted TME Hallmarks
F
−1
0
1
Spatial Dependency
G
−1.0
−0.5
0.0
0.5
1.0
Spatial Dependency
H
Spatial Dependency
0.4
0.2
0
−0.2
−0.4
TME Predictors
Predicted Neoplastic Hallmarks
I
Spatial Dependency
0.4
0.2
0
−0.2
Neoplastic Predictors
Predicted TME Hallmarks
Pediatric TME Hallmarks
Sustaining Proliferative Signaling
Unlocking Phenotypic Plasticity
Avoiding Immune Destruction
Tumor−Promoting Inflammation
Activating Invasion and Metastasis
Inducing Angiogenesis
Stochastic Hallmarks
Senescent Cells
Deregulating Cellular Energetics
Evading Growth Suppressors
Resisting Cell Death
Pediatric Neoplastic
Nonmutational Epigenetic Reprogramming
Enabling Replicative Immortality
Genome Instability and Mutation

Figure S4

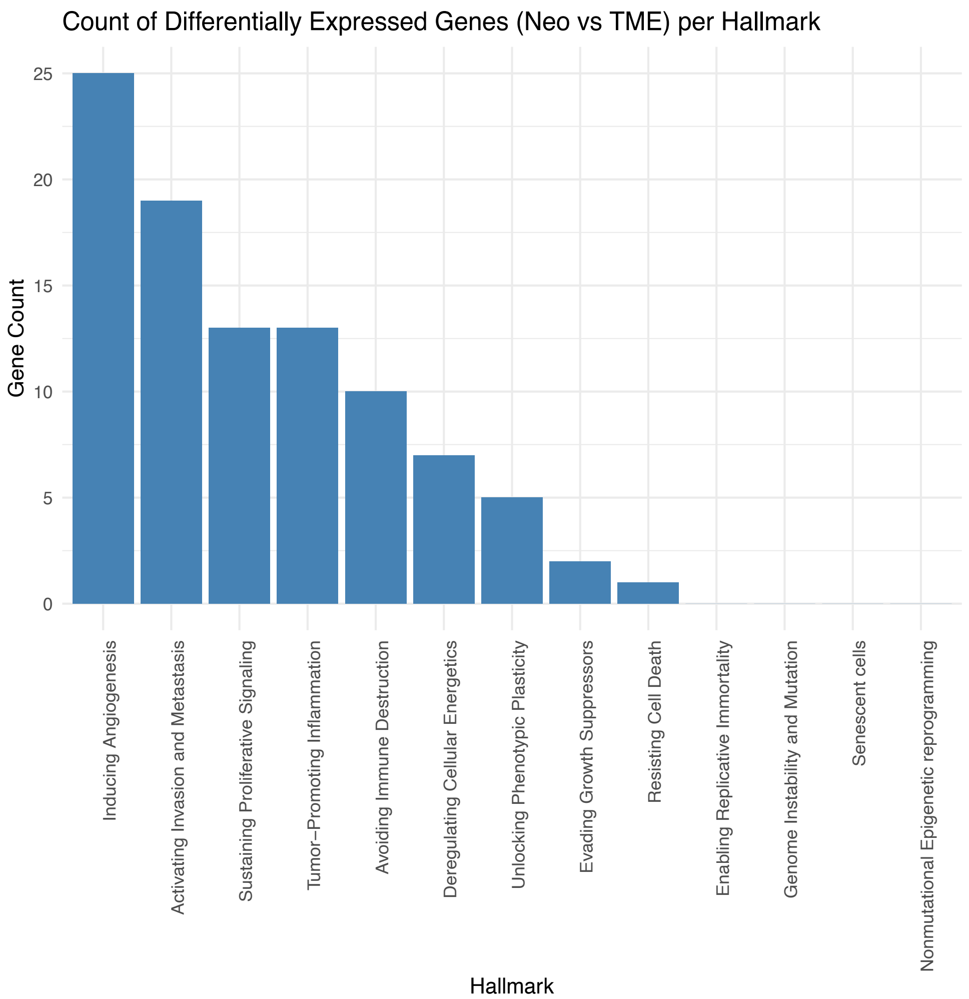

Count of Differentially Expressed Genes (Neo vs TME) per Hallmark
Gene Count
25
20
15
10
5
0
Inducing Angiogenesis
Activating Invasion and Metastasis
Sustaining Proliferative Signaling
Tumor−Promoting Inflammation
Avoiding Immune Destruction
Deregulating Cellular Energetics
Unlocking Phenotypic Plasticity
Evading Growth Suppressors
Resisting Cell Death
Enabling Replicative Immortality
Genome Instability and Mutation
Senescent cells
Nonmutational Epigenetic reprogramming
Hallmark


Figure S5

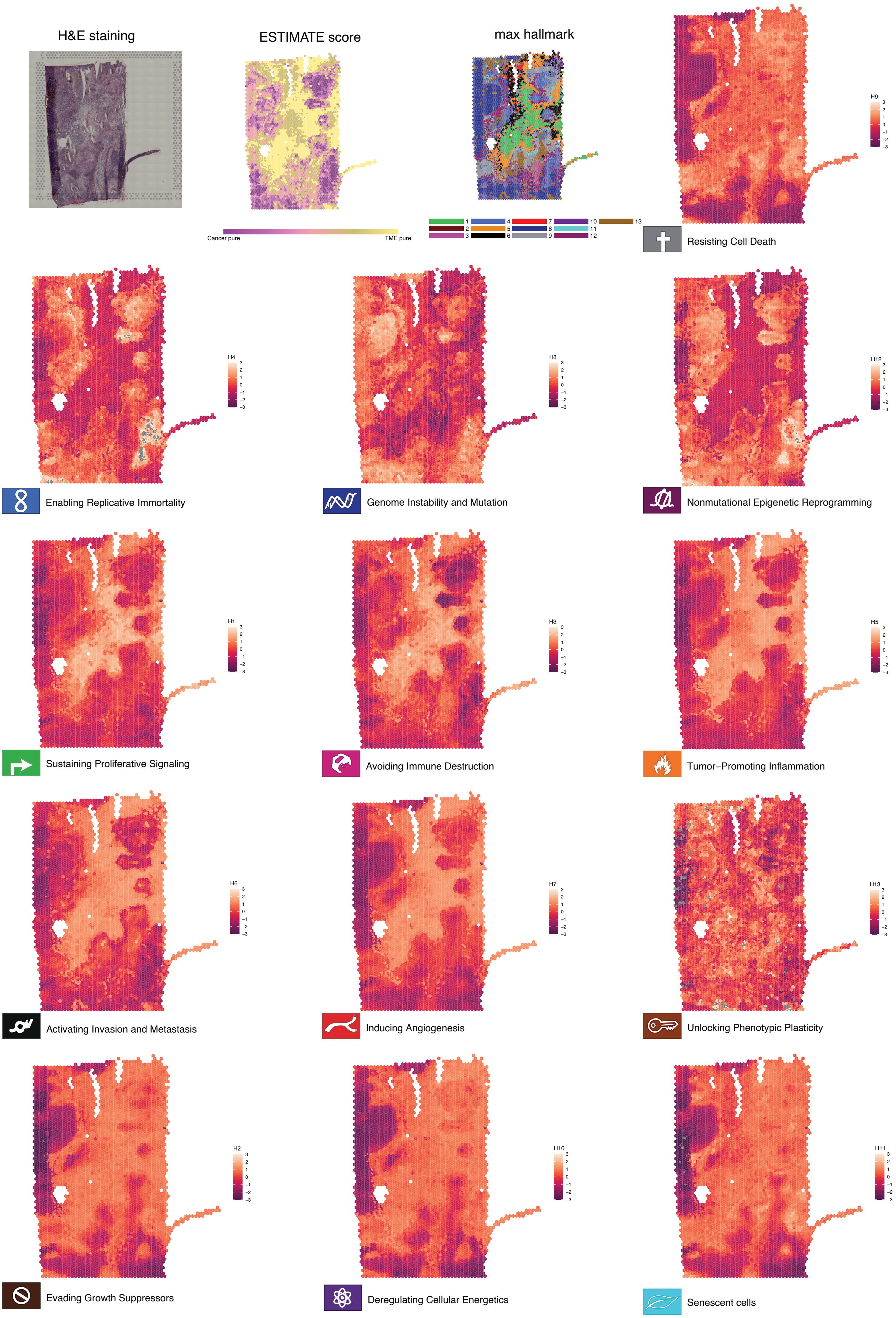
H&E staining
ESTIMATE score
Cancer pure
TME pure
max hallmark
1
2
3
4
5
6
7
8
9
10
11
12
13
H9
Resisting Cell Death
H4
Enabling Replicative Immortality
H8
Genome Instability and Mutation
H12
Nonmutational Epigenetic Reprogramming
H1
Sustaining Proliferative Signaling
H3
Avoiding Immune Destruction
H5
Tumor−Promoting Inflammation
H6
Activating Invasion and Metastasis
H7
Inducing Angiogenesis
H13
Unlocking Phenotypic Plasticity
H2
Evading Growth Suppressors
H10
Deregulating Cellular Energetics
H11
Senescent cells

Figure S6

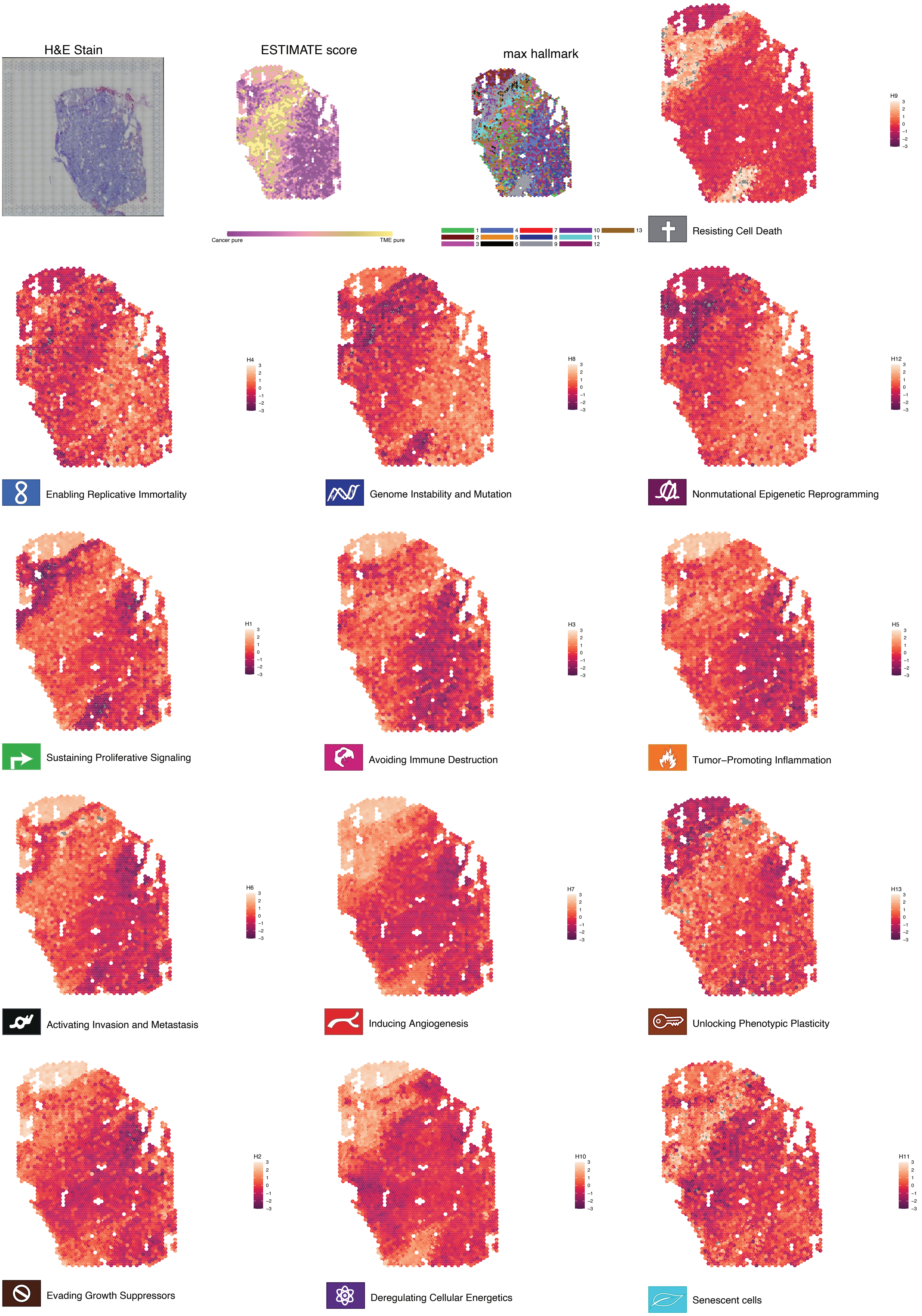
H&E Stain
ESTIMATE score
Cancer pure
TME pure
max hallmark
1
2
3
4
5
6
7
8
9
10
11
12
13
H9
Resisting Cell Death
H4
Enabling Replicative Immortality
H8
Genome Instability and Mutation
H12
Nonmutational Epigenetic Reprogramming
H1
Sustaining Proliferative Signaling
H3
Avoiding Immune Destruction
H5
Tumor−Promoting Inflammation
H6
Activating Invasion and Metastasis
H7
Inducing Angiogenesis
H13
Unlocking Phenotypic Plasticity
H2
Evading Growth Suppressors
H10
Deregulating Cellular Energetics
H11
Senescent cells

Figure S7

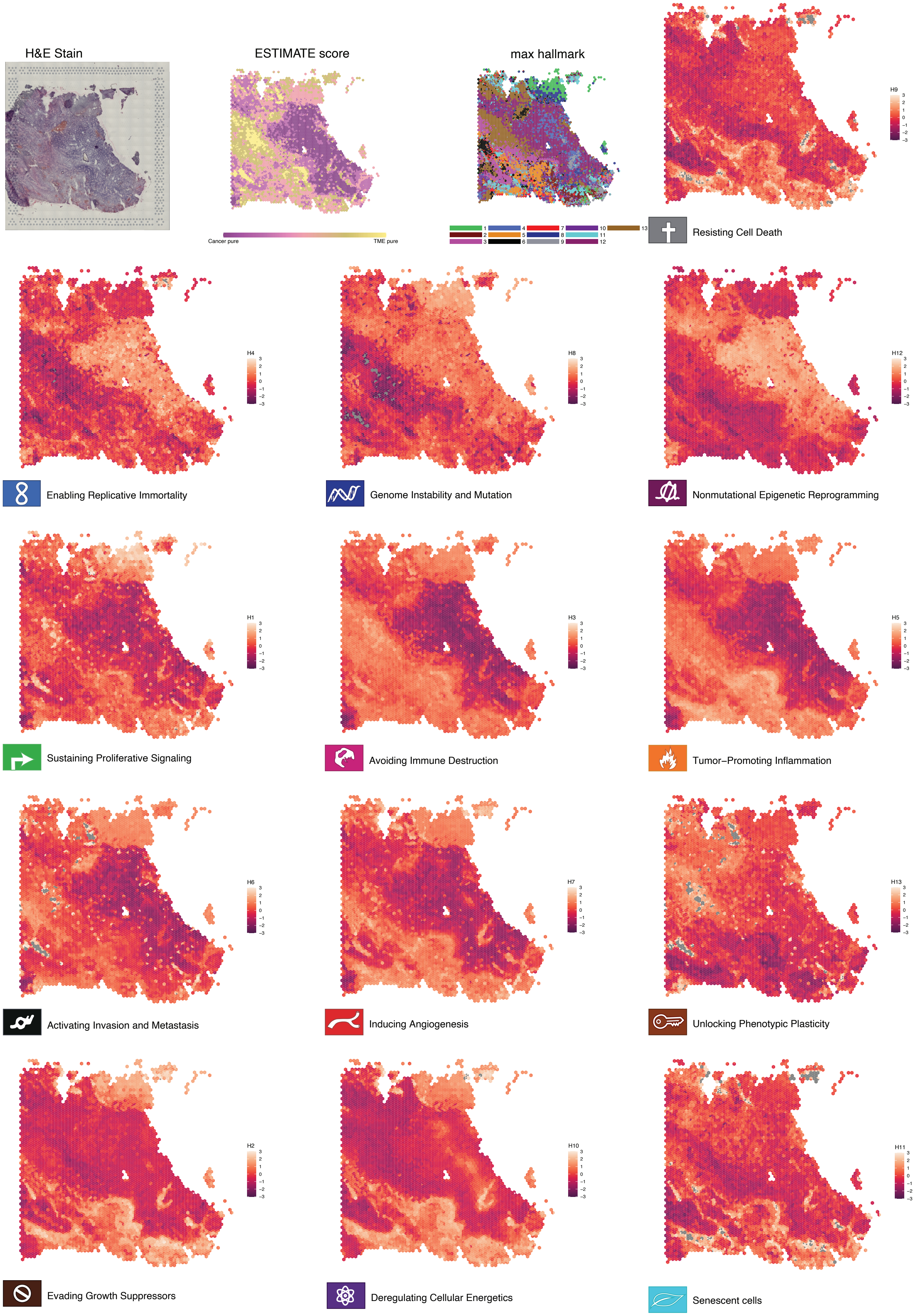
H&E Stain
ESTIMATE score
Cancer pure
TME pure
max hallmark
1
2
3
4
5
6
7
8
9
10
11
12
13
H9
Resisting Cell Death
H4
Enabling Replicative Immortality
H8
Genome Instability and Mutation
H12
Nonmutational Epigenetic Reprogramming
H1
Sustaining Proliferative Signaling
H3
Avoiding Immune Destruction
H5
Tumor−Promoting Inflammation
H6
Activating Invasion and Metastasis
H7
Inducing Angiogenesis
H13
Unlocking Phenotypic Plasticity
H2
Evading Growth Suppressors
H10
Deregulating Cellular Energetics
H11
Senescent cells

Figure S8

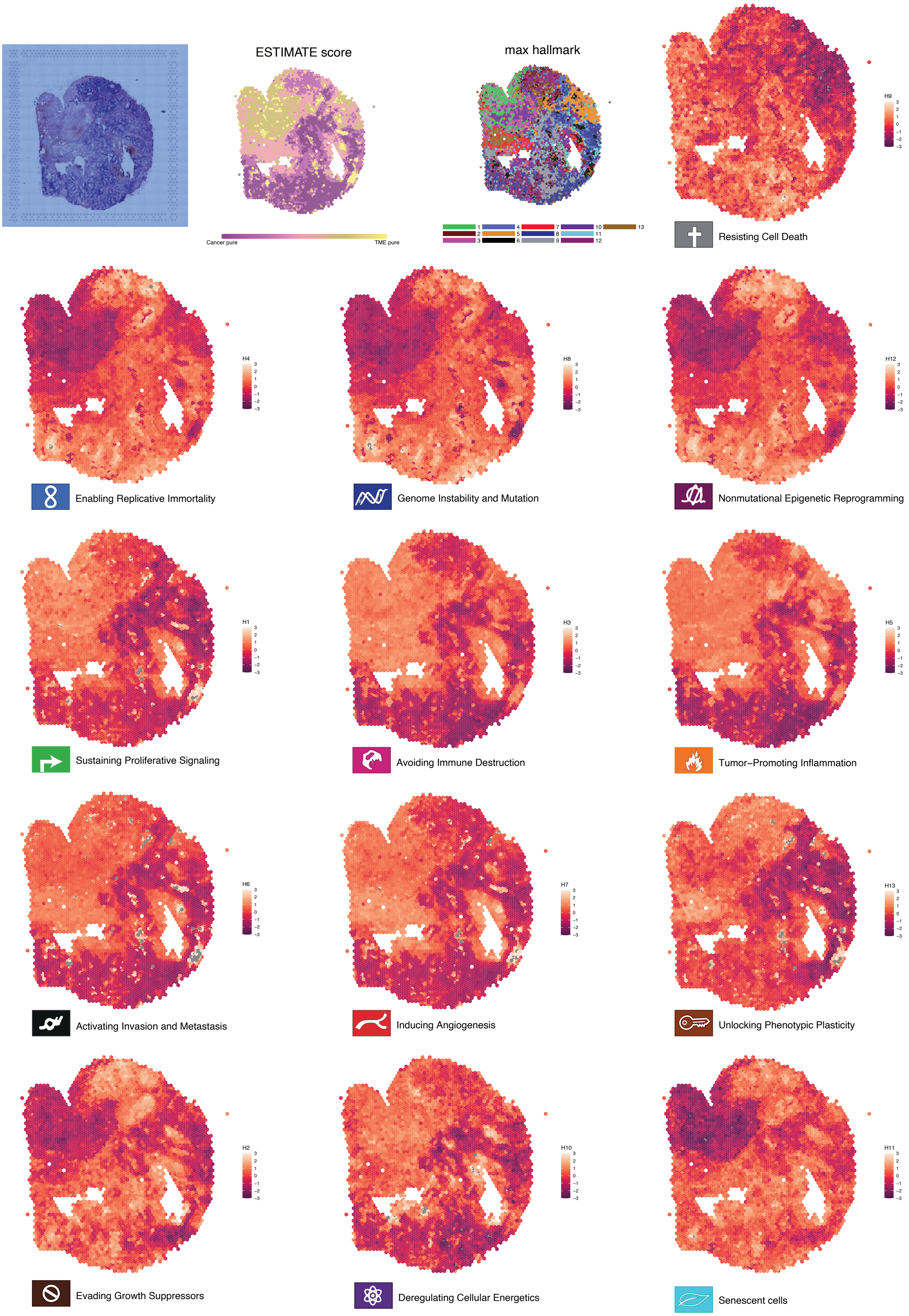
ESTIMATE score
Cancer pure
TME pure
max hallmark
Resisting Cell Death
Enabling Replicative Immortality
Genome Instability and Mutation
Nonmutational Epigenetic Reprogramming
Sustaining Proliferative Signaling
Avoiding Immune Destruction
Tumor–Promoting Inflammation
Activating Invasion and Metastasis
Inducing Angiogenesis
Unlocking Phenotypic Plasticity
Evading Growth Suppressors
Deregulating Cellular Energetics
Senescent cells

Figure S9

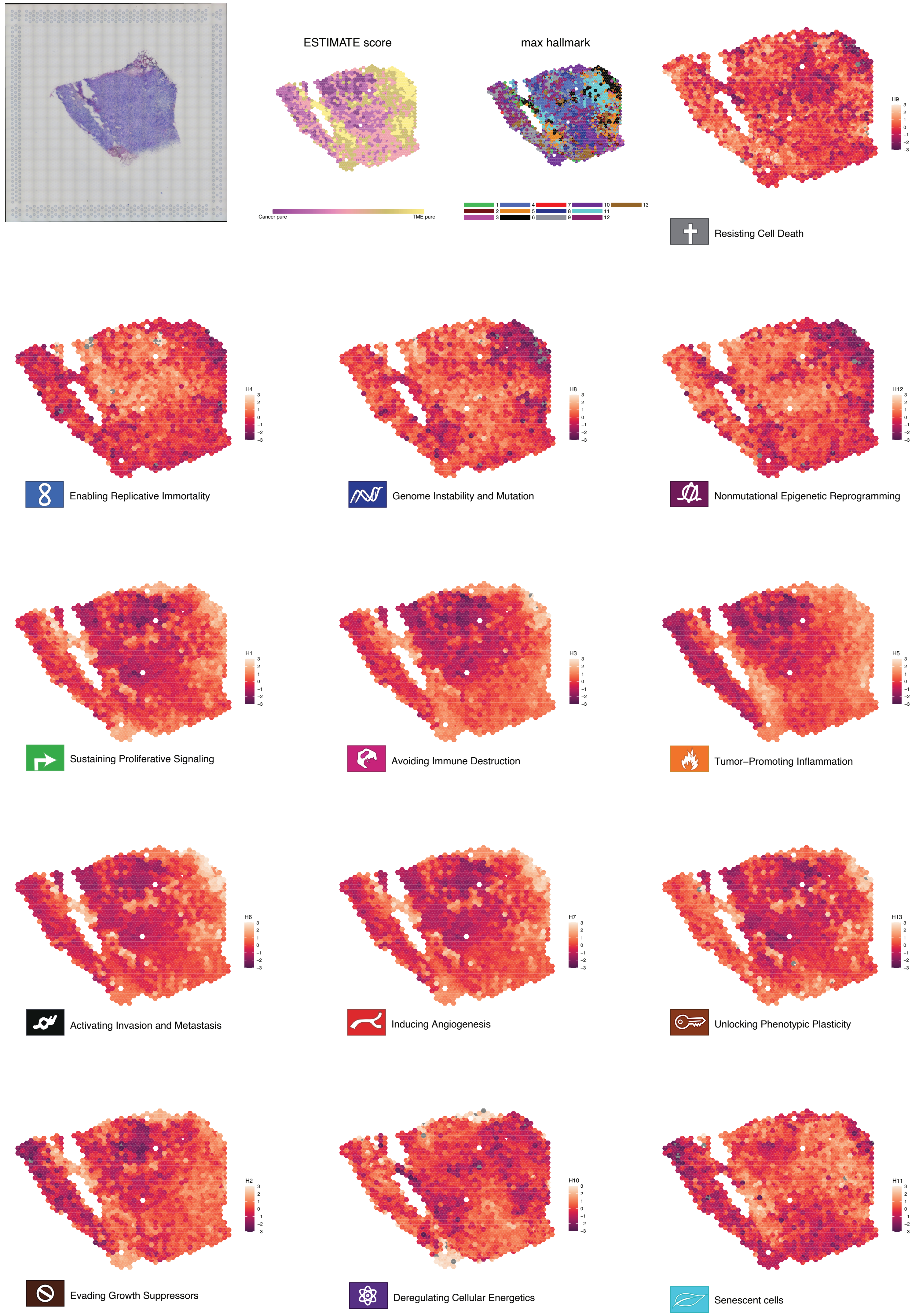
ESTIMATE score
Cancer pure
TME pure
max hallmark
1 2 3 4 5 6 7 8 9 10 11 12 13
H9
Resisting Cell Death
H4
Enabling Replicative Immortality
H8
Genome Instability and Mutation
H12
Nonmutational Epigenetic Reprogramming
H1
Sustaining Proliferative Signaling
H3
Avoiding Immune Destruction
H5
Tumor–Promoting Inflammation
H6
Activating Invasion and Metastasis
H7
Inducing Angiogenesis
H13
Unlocking Phenotypic Plasticity
H2
Evading Growth Suppressors
H10
Deregulating Cellular Energetics
H11
Senescent cells

Figure S10

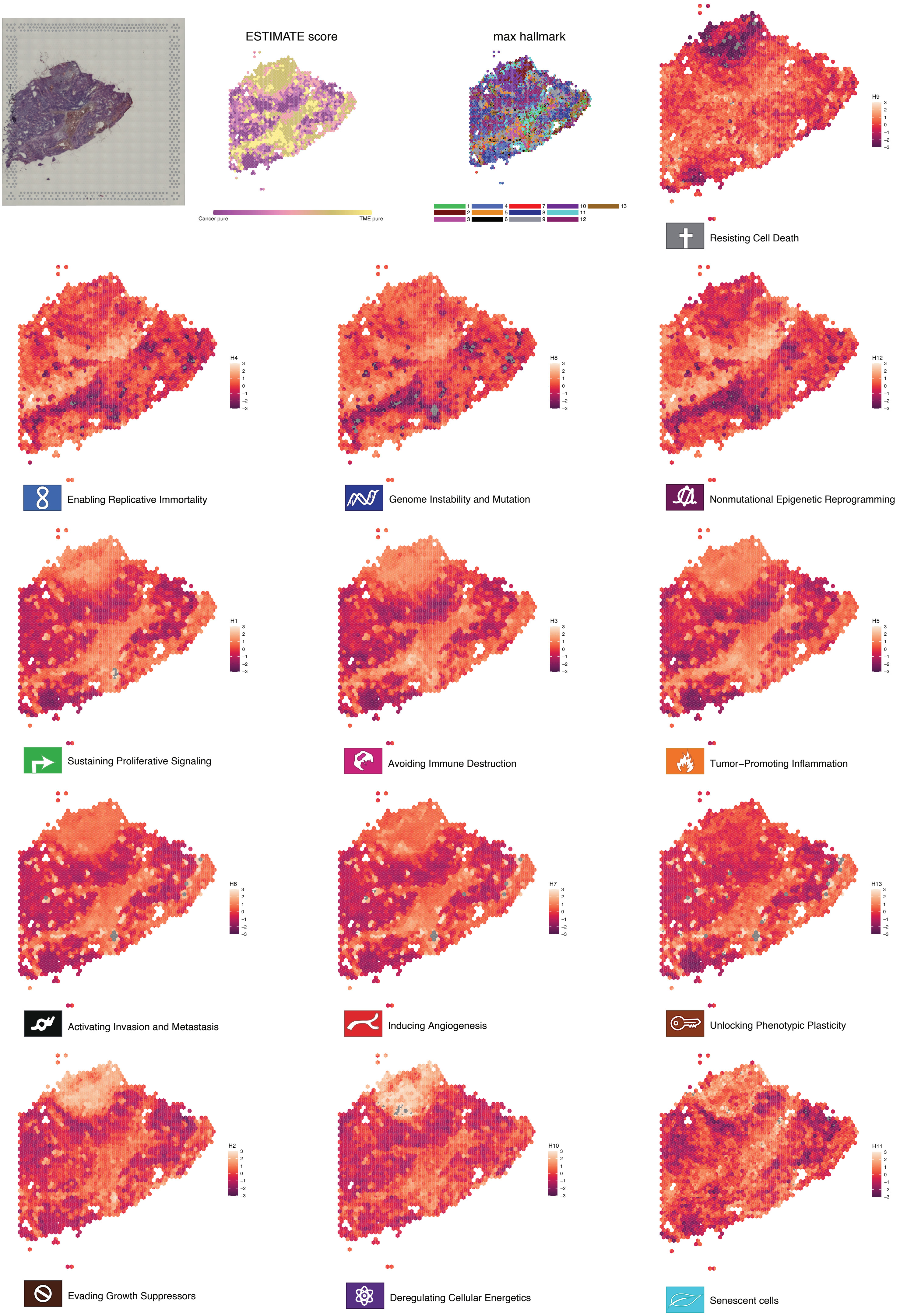
ESTIMATE score
Cancer pure
TME pure
max hallmark
1 2 3 4 5 6 7 8 9 10 11 12 13
H9
Resisting Cell Death
H4
Enabling Replicative Immortality
H8
Genome Instability and Mutation
H12
Nonmutational Epigenetic Reprogramming
H1
Sustaining Proliferative Signaling
H3
Avoiding Immune Destruction
H5
Tumor−Promoting Inflammation
H6
Activating Invasion and Metastasis
H7
Inducing Angiogenesis
H13
Unlocking Phenotypic Plasticity
H2
Evading Growth Suppressors
H10
Deregulating Cellular Energetics
H11
Senescent cells

Figure S11

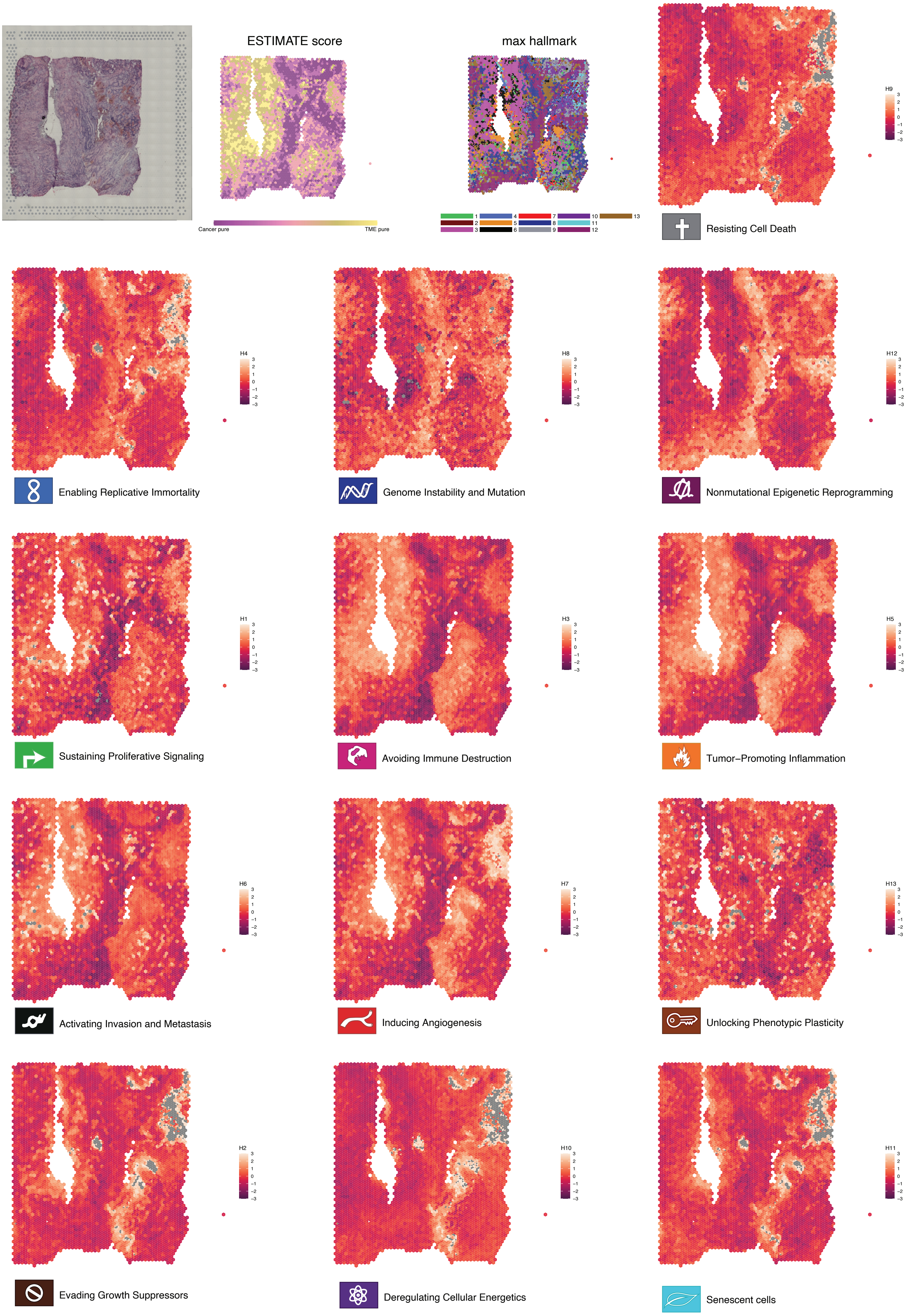
ESTIMATE score
Cancer pure
TME pure
max hallmark
Resisting Cell Death
H9
Enabling Replicative Immortality
H4
Genome Instability and Mutation
H8
Nonmutational Epigenetic Reprogramming
H12
Sustaining Proliferative Signaling
H1
Avoiding Immune Destruction
H3
Tumor−Promoting Inflammation
H5
Activating Invasion and Metastasis
H6
Inducing Angiogenesis
H7
Unlocking Phenotypic Plasticity
H13
Evading Growth Suppressors
H2
Deregulating Cellular Energetics
H10
Senescent cells
H11

Figure S12

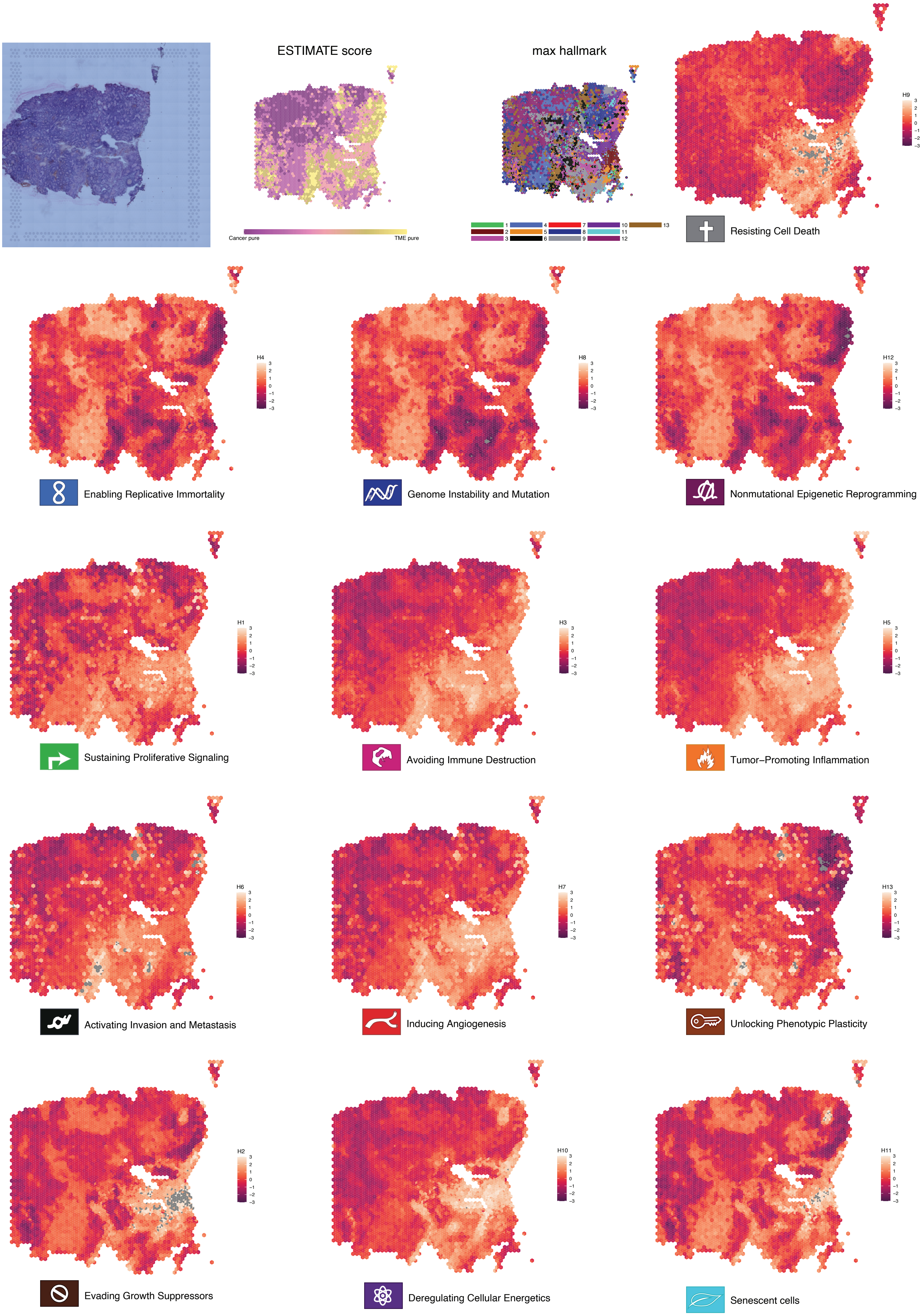

ESTIMATE score
Cancer pure
TME pure
max hallmark
1
2
3
4
5
6
7
8
9
10
11
12
13
Resisting Cell Death
H9
Enabling Replicative Immortality
H4
Genome Instability and Mutation
H8
Nonmutational Epigenetic Reprogramming
H12
Sustaining Proliferative Signaling
H1
Avoiding Immune Destruction
H3
Tumor–Promoting Inflammation
H5
Activating Invasion and Metastasis
H6
Inducing Angiogenesis
H7
Unlocking Phenotypic Plasticity
H13
Evading Growth Suppressors
H2
Deregulating Cellular Energetics
H10
Senescent cells
H11


Figure S13

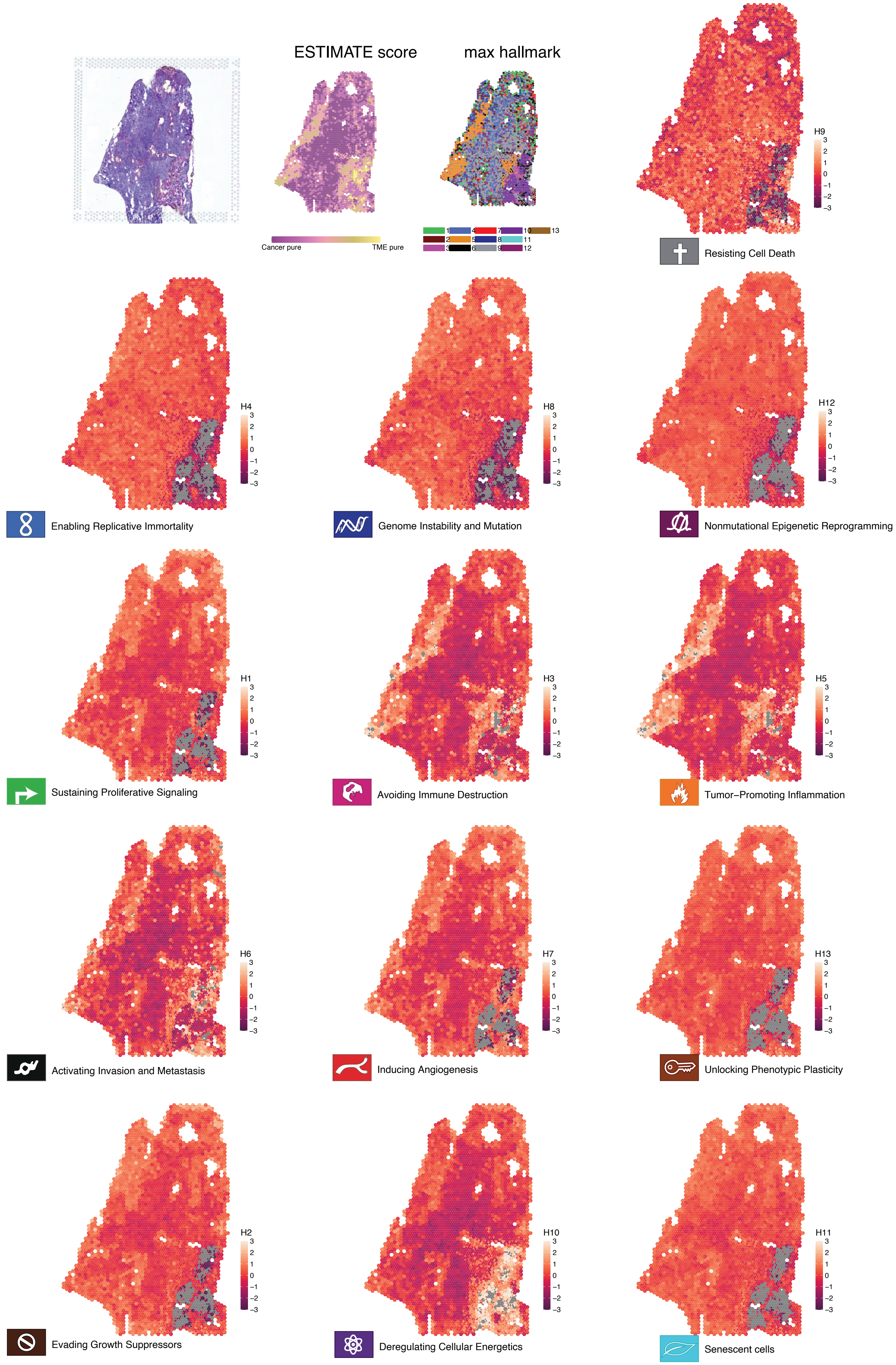

ESTIMATE score
max hallmark
Cancer pure
TME pure
1
2
3
4
5
6
7
8
9
10
11
12
13
H9
Resisting Cell Death
H4
Enabling Replicative Immortality
H8
Genome Instability and Mutation
H12
Nonmutational Epigenetic Reprogramming
H1
Sustaining Proliferative Signaling
H3
Avoiding Immune Destruction
H5
Tumor–Promoting Inflammation
H6
Activating Invasion and Metastasis
H7
Inducing Angiogenesis
H13
Unlocking Phenotypic Plasticity
H2
Evading Growth Suppressors
H10
Deregulating Cellular Energetics
H11
Senescent cells


Figure S14

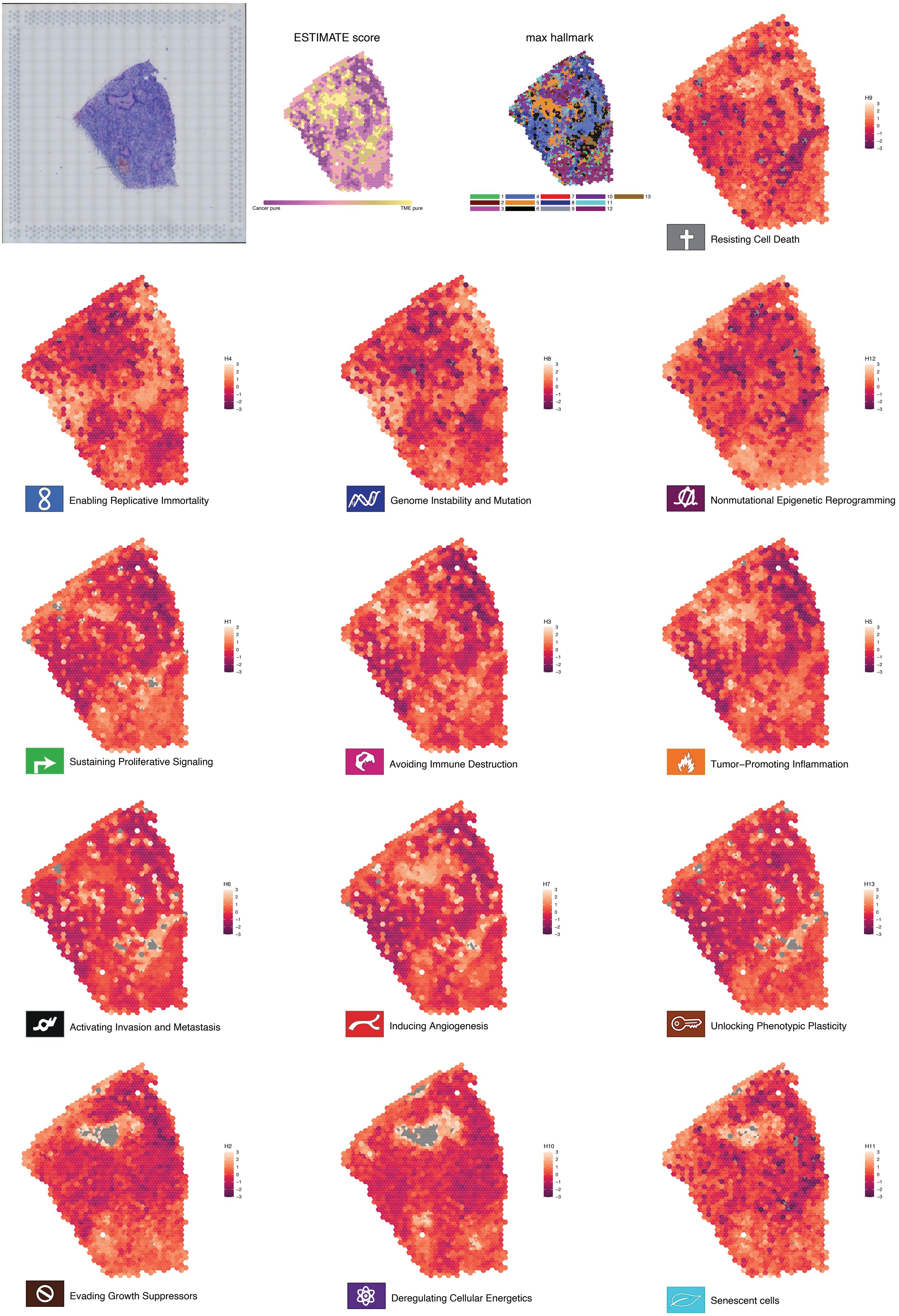
ESTIMATE score
Cancer pure
TME pure
max hallmark
Resisting Cell Death
Enabling Replicative Immortality
Genome Instability and Mutation
Nonmutational Epigenetic Reprogramming
Sustaining Proliferative Signaling
Avoiding Immune Destruction
Tumor−Promoting Inflammation
Activating Invasion and Metastasis
Inducing Angiogenesis
Unlocking Phenotypic Plasticity
Evading Growth Suppressors
Deregulating Cellular Energetics
Senescent cells

Figure S15

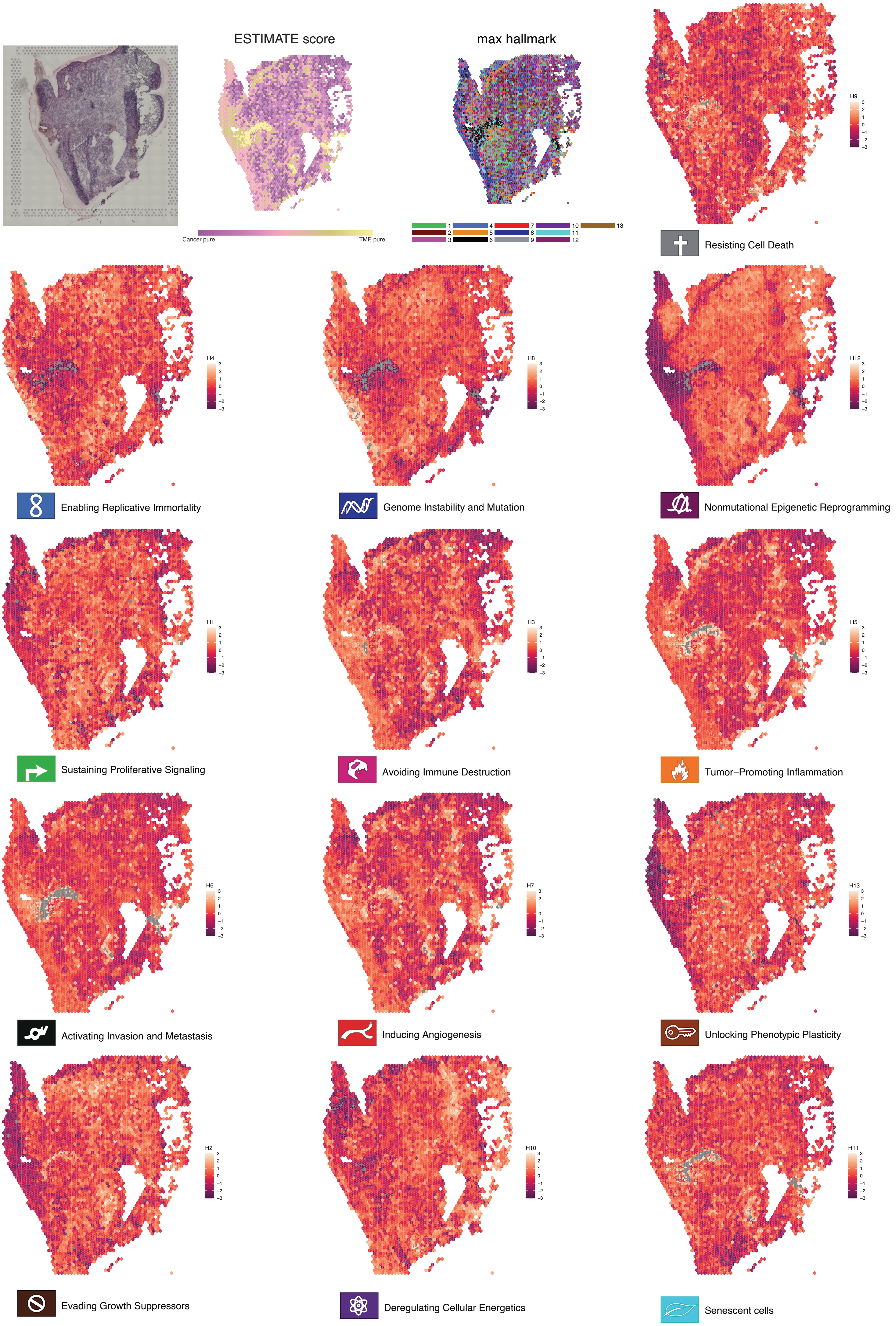
ESTIMATE score
Cancer pure
TME pure
max hallmark
1
2
3
4
5
6
7
8
9
10
11
12
13
H9
Resisting Cell Death
H4
Enabling Replicative Immortality
H8
Genome Instability and Mutation
H12
Nonmutational Epigenetic Reprogramming
H1
Sustaining Proliferative Signaling
H3
Avoiding Immune Destruction
H5
Tumor−Promoting Inflammation
H6
Activating Invasion and Metastasis
H7
Inducing Angiogenesis
H13
Unlocking Phenotypic Plasticity
H2
Evading Growth Suppressors
H10
Deregulating Cellular Energetics
H11
Senescent cells

Figure S16

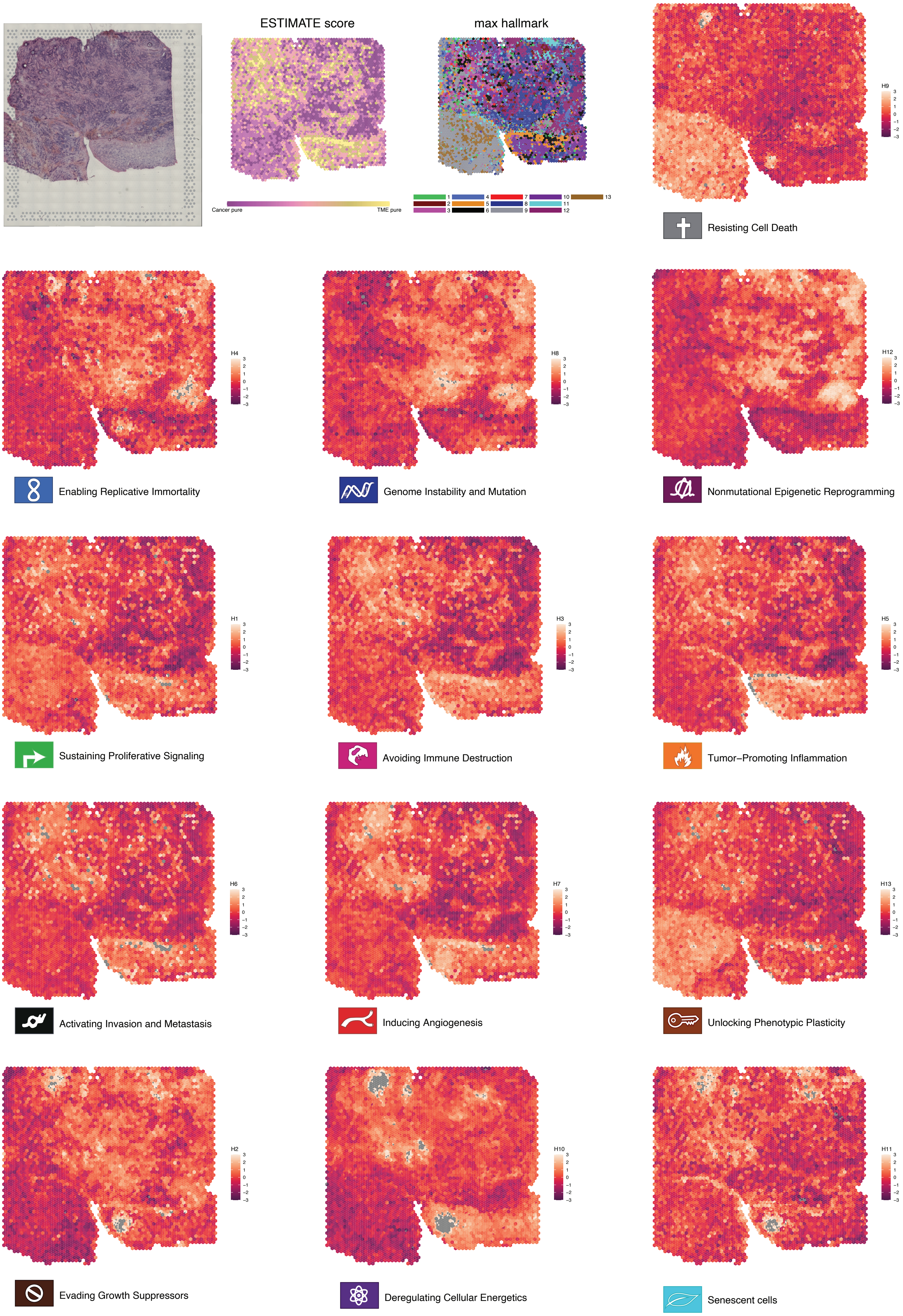
ESTIMATE score
Cancer pure
TME pure
max hallmark
1
2
3
4
5
6
7
8
9
10
11
12
13
H9
Resisting Cell Death
H4
Enabling Replicative Immortality
H8
Genome Instability and Mutation
H12
Nonmutational Epigenetic Reprogramming
H1
Sustaining Proliferative Signaling
H3
Avoiding Immune Destruction
H5
Tumor−Promoting Inflammation
H6
Activating Invasion and Metastasis
H7
Inducing Angiogenesis
H13
Unlocking Phenotypic Plasticity
H2
Evading Growth Suppressors
H10
Deregulating Cellular Energetics
H11
Senescent cells

Figure S17

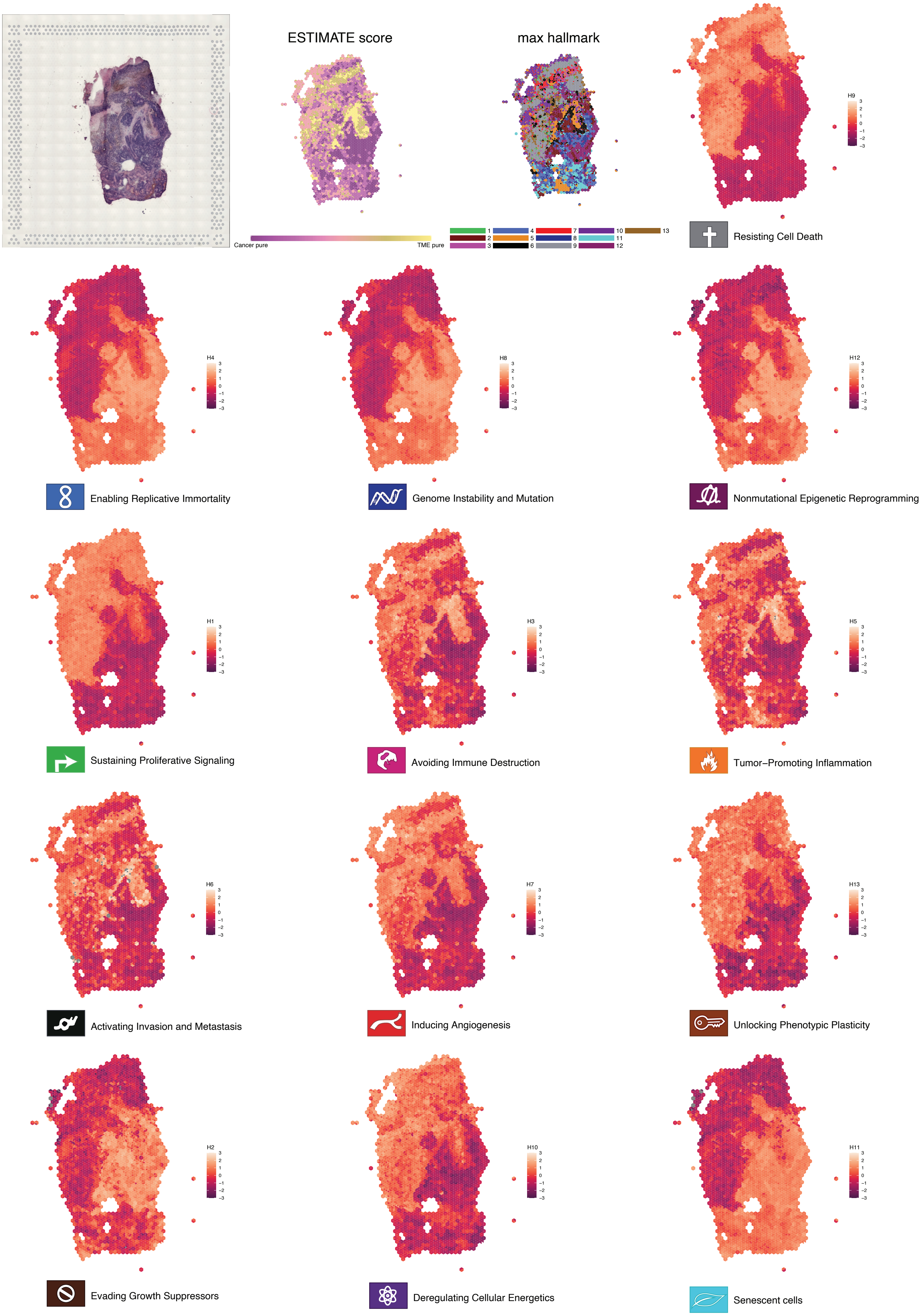

ESTIMATE score
Cancer pure
TME pure
max hallmark
1
2
3
4
5
6
7
8
9
10
11
12
13
H9
3
2
1
0
−1
−2
−3
Resisting Cell Death
H4
Enabling Replicative Immortality
H8
Genome Instability and Mutation
H12
Nonmutational Epigenetic Reprogramming
H1
Sustaining Proliferative Signaling
H3
Avoiding Immune Destruction
H5
Tumor−Promoting Inflammation
H6
Activating Invasion and Metastasis
H7
Inducing Angiogenesis
H13
Unlocking Phenotypic Plasticity
H2
Evading Growth Suppressors
H10
Deregulating Cellular Energetics
H11
Senescent cells


Figure S18

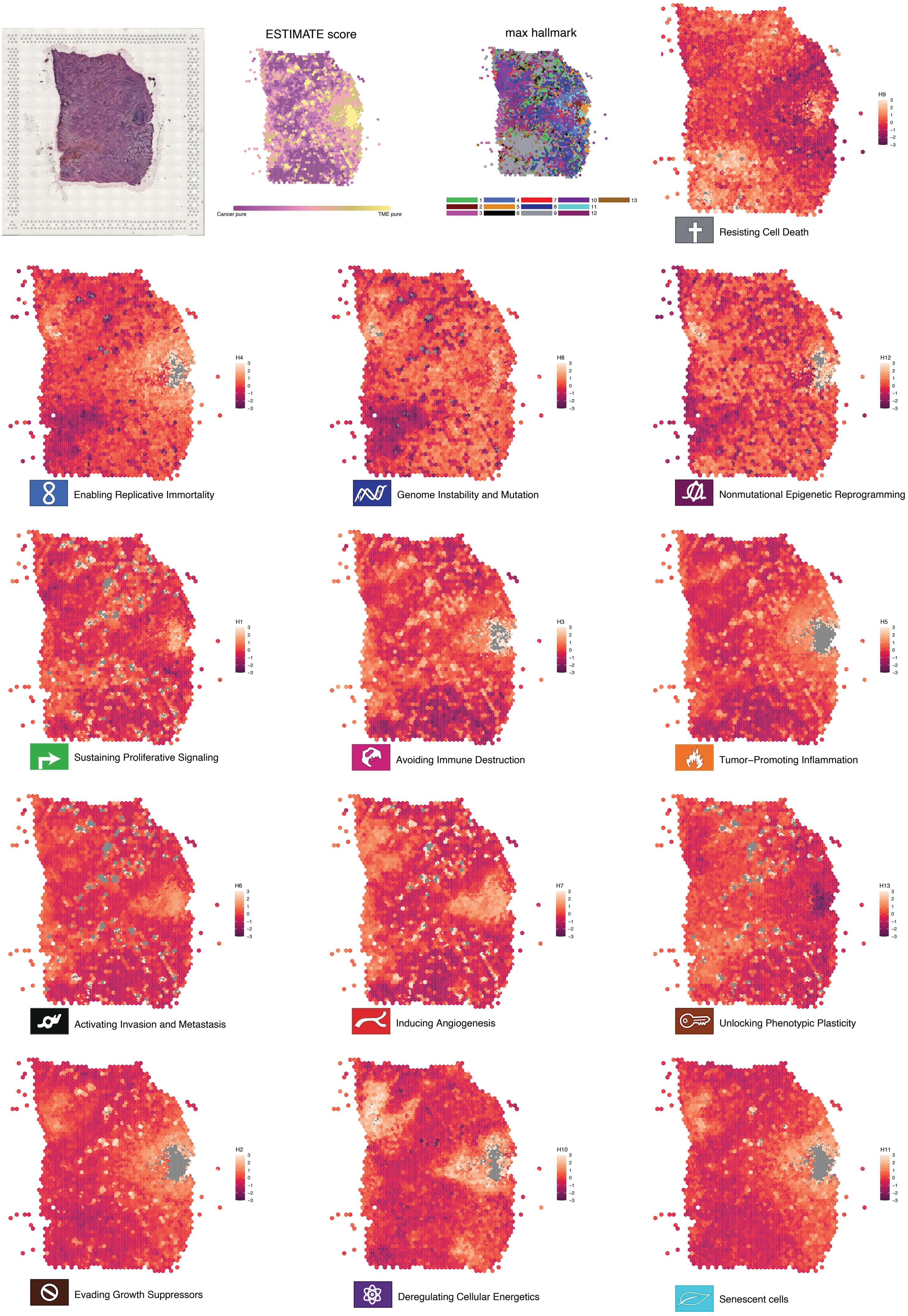
ESTIMATE score
Cancer pure
TME pure
max hallmark
1
2
3
4
5
6
7
8
9
10
11
12
13
H9
Resisting Cell Death
H4
Enabling Replicative Immortality
H8
Genome Instability and Mutation
H12
Nonmutational Epigenetic Reprogramming
H1
Sustaining Proliferative Signaling
H3
Avoiding Immune Destruction
H5
Tumor−Promoting Inflammation
H6
Activating Invasion and Metastasis
H7
Inducing Angiogenesis
H13
Unlocking Phenotypic Plasticity
H2
Evading Growth Suppressors
H10
Deregulating Cellular Energetics
H11
Senescent cells

Figure S19

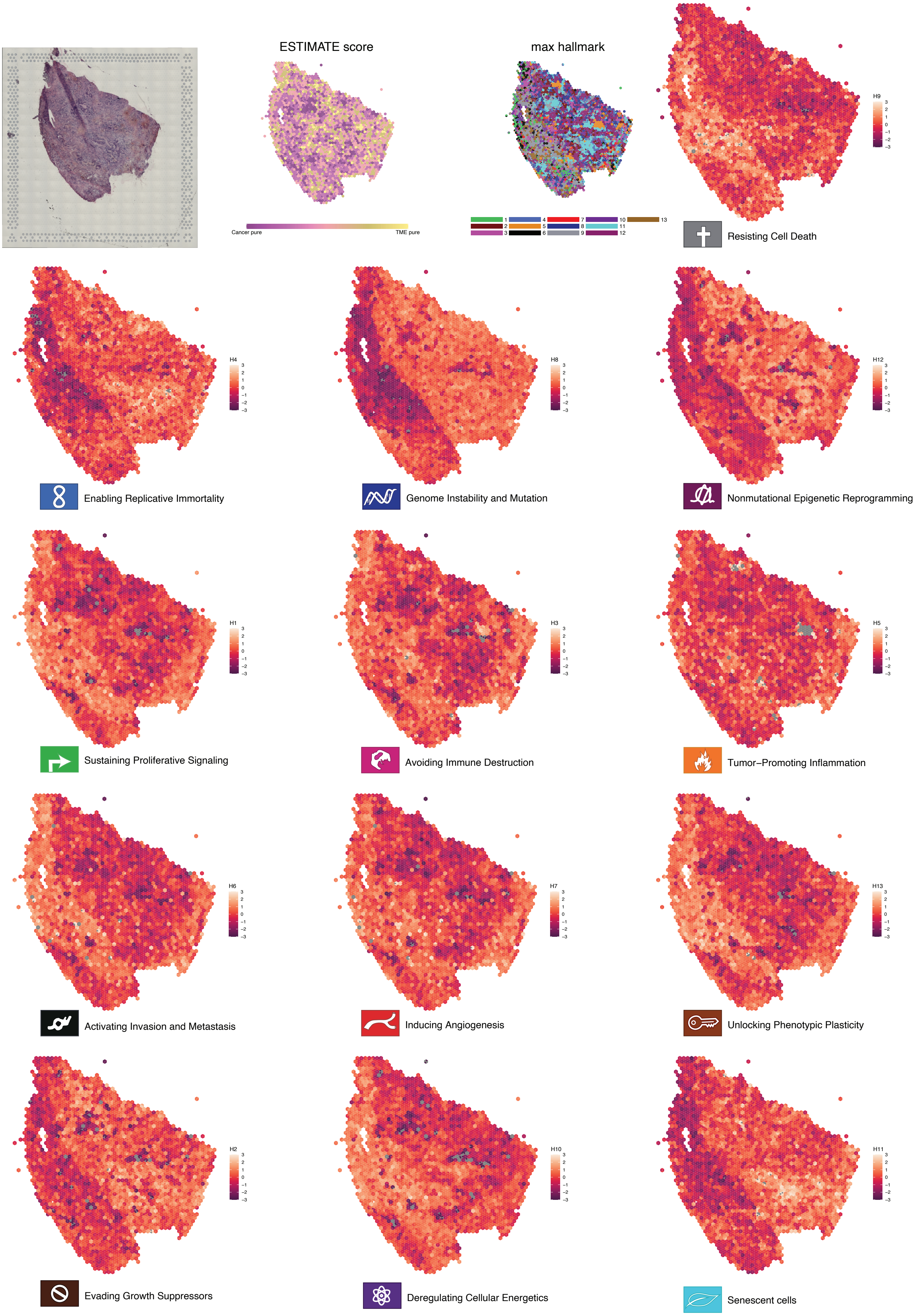
ESTIMATE score
Cancer pure
TME pure
max hallmark
1
2
3
4
5
6
7
8
9
10
11
12
13
H9
Resisting Cell Death
H4
Enabling Replicative Immortality
H8
Genome Instability and Mutation
H12
Nonmutational Epigenetic Reprogramming
H1
Sustaining Proliferative Signaling
H3
Avoiding Immune Destruction
H5
Tumor−Promoting Inflammation
H6
Activating Invasion and Metastasis
H7
Inducing Angiogenesis
H13
Unlocking Phenotypic Plasticity
H2
Evading Growth Suppressors
H10
Deregulating Cellular Energetics
H11
Senescent cells

Figure S20

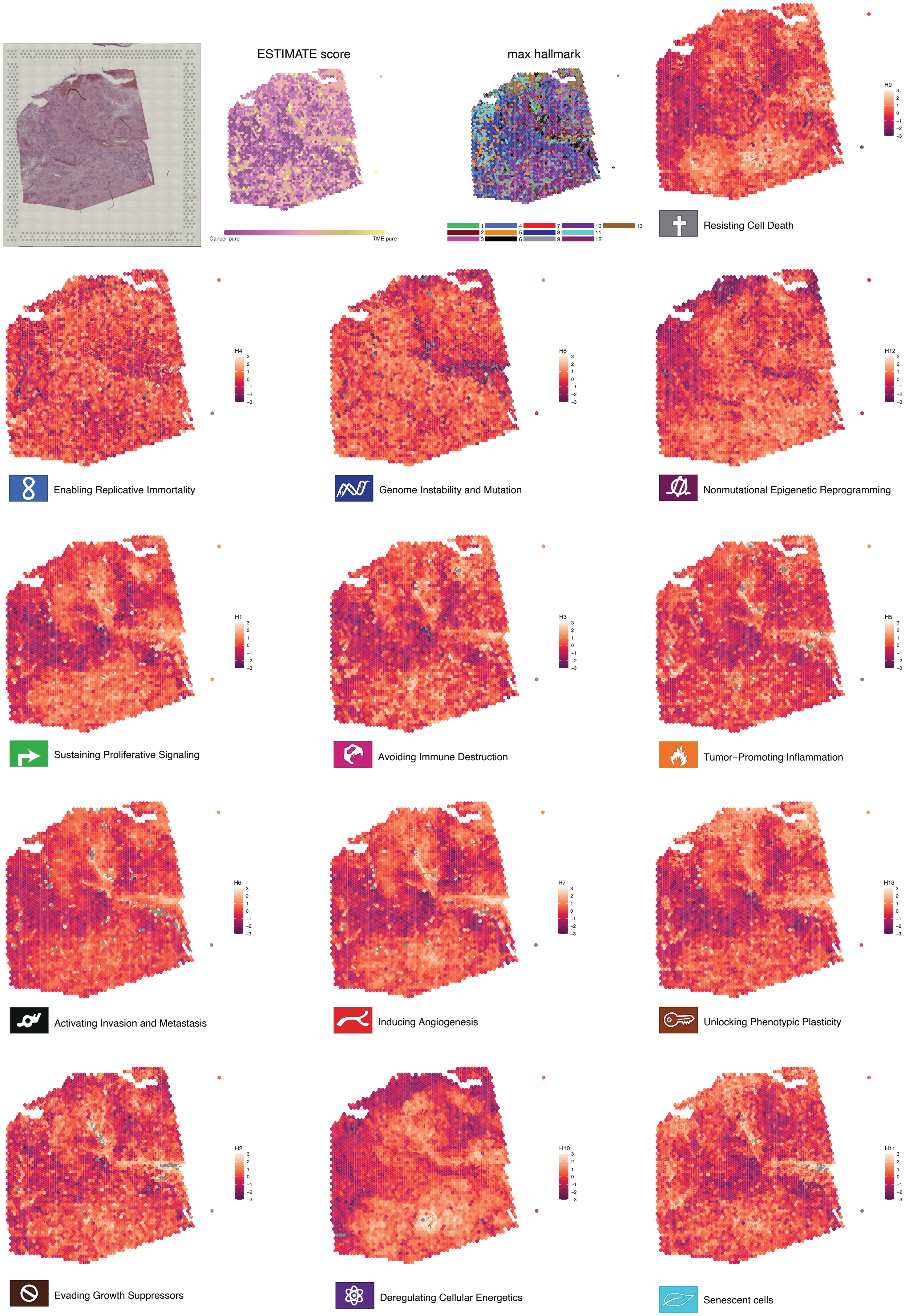
ESTIMATE score
Cancer pure
TME pure
max hallmark
1 2 3 4 5 6 7 8 9 10 11 12 13
H9
3 2 1 0 −1 −2 −3
Resisting Cell Death
H4
Enabling Replicative Immortality
H8
Genome Instability and Mutation
H12
Nonmutational Epigenetic Reprogramming
H1
Sustaining Proliferative Signaling
H3
Avoiding Immune Destruction
H5
Tumor−Promoting Inflammation
H6
Activating Invasion and Metastasis
H7
Inducing Angiogenesis
H13
Unlocking Phenotypic Plasticity
H2
Evading Growth Suppressors
H10
Deregulating Cellular Energetics
H11
Senescent cells

Figure S21

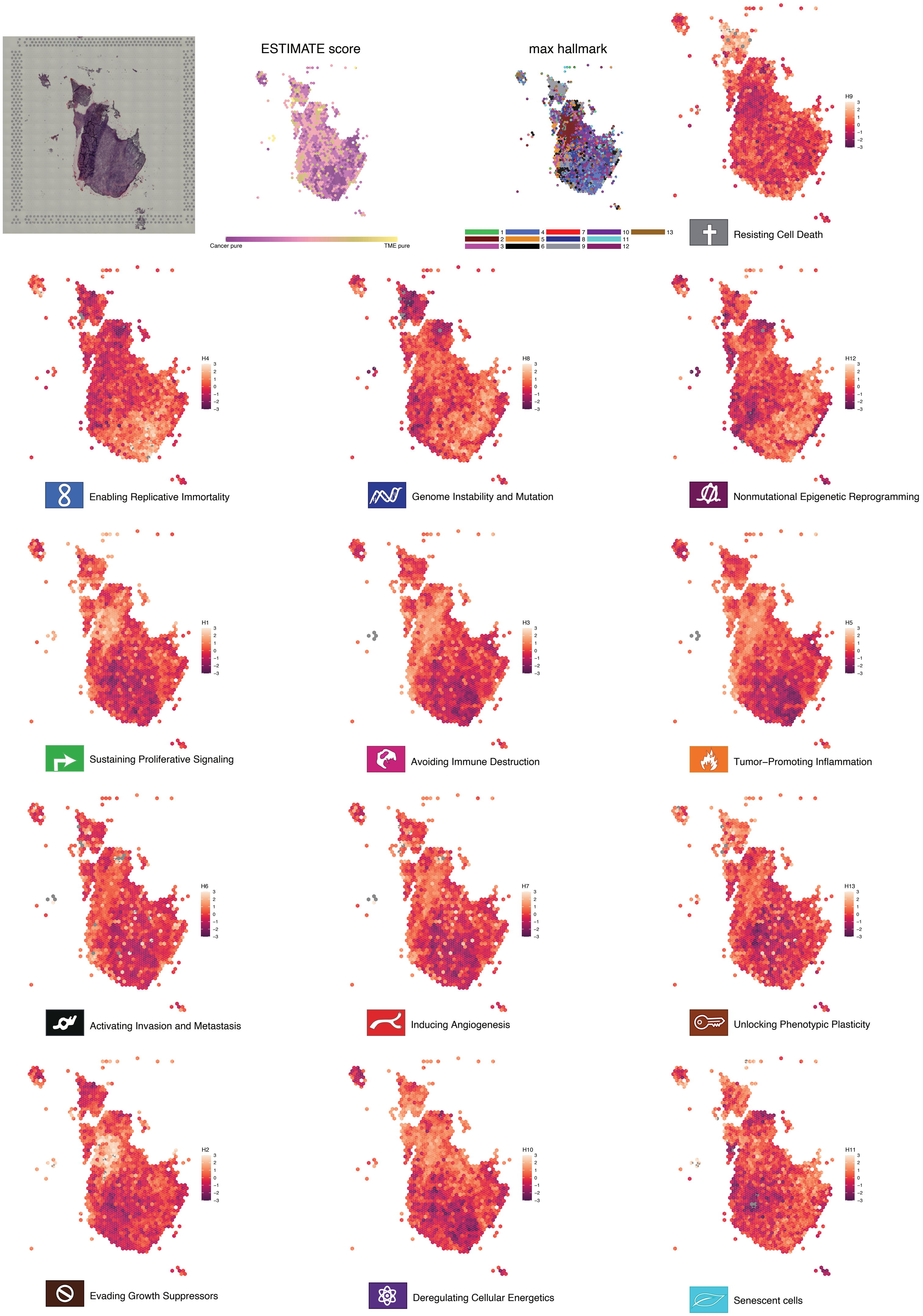
ESTIMATE score
Cancer pure
TME pure
max hallmark
1
2
3
4
5
6
7
8
9
10
11
12
13
H9
Resisting Cell Death
H4
Enabling Replicative Immortality
H8
Genome Instability and Mutation
H12
Nonmutational Epigenetic Reprogramming
H1
Sustaining Proliferative Signaling
H3
Avoiding Immune Destruction
H5
Tumor–Promoting Inflammation
H6
Activating Invasion and Metastasis
H7
Inducing Angiogenesis
H13
Unlocking Phenotypic Plasticity
H2
Evading Growth Suppressors
H10
Deregulating Cellular Energetics
H11
Senescent cells

Figure S22
